# Multi-layered chromatin proteomics identifies cell vulnerabilities in DNA repair

**DOI:** 10.1101/2021.08.05.455226

**Authors:** Gianluca Sigismondo, Lavinia Arseni, Nicolàs Palacio-Escat, Thomas G Hofmann, Martina Seiffert, Jeroen Krijgsveld

## Abstract

The DNA damage response (DDR) is essential to maintain genome stability, and its deregulation predisposes to carcinogenesis while encompassing attractive targets for cancer therapy. Chromatin governs the DDR *via* the concerted interplay among different layers, including DNA, histone post-translational modifications (hPTMs), and chromatin-associated proteins. Here we employ multi-layered proteomics to characterize chromatin-mediated functional interactions of repair proteins, signatures of hPTMs, and the DNA-bound proteome during DNA double-strand break (DSB) repair at high temporal resolution. Our data illuminate the dynamics of known and novel DDR-associated factors both at chromatin and at DSBs. We functionally attribute novel chromatin-associated proteins to repair by non-homologous end-joining (NHEJ), homologous recombination (HR) and DSB repair pathway choice. We reveal histone reader ATAD2, microtubule organizer TPX2 and histone methyltransferase G9A as regulators of HR and PARP-inhibitor sensitivity. Furthermore, we distinguish hPTMs that are globally induced by DNA damage from those specifically acquired at sites flanking DSBs (γH2AX *foci*-specific), and profiled their dynamics during the DDR. Integration of complementary chromatin layers implicates G9A-mediated monomethylation of H3K56 at DSBs in HR. Our data provide a dynamic chromatin-centered view of the DDR that can be further mined to identify novel mechanistic links and cell vulnerabilities in DSB repair.

## INTRODUCTION

DNA damage represents a major risk for genome stability, and among the different types of lesions, double-strand breaks (DSBs) are the most detrimental; indeed if not properly repaired these lesions predispose to DNA mutations and loss of genomic information. To prevent genome instability, two main repair mechanisms have evolved: the error-prone non-homologous end joining (NHEJ) based on the fast ligation of DSBs, and the homologous-directed recombination (HDR or HR), where sister chromatid is used as template for error-free DSB repair (1-4). Notably, the DNA damage repair occurs in the context of chromatin, a higher ordered structure composed of DNA wrapped around histone proteins, and stabilized by non-histone components (5). Upon DSB formation, chromatin determinants reorganize the structure surrounding the lesion, activate a specific signaling cascade, and recruit the repair machinery for efficient DSB resolution *via* either NHEJ or HR (6). In particular, the sensor complex composed of MRE11, RAD50 and NBN (MRN complex) rapidly accumulates at damaged sites where it promotes the ATM-mediated phosphorylation of the histone variant H2A.X (referred to as γH2AX). This histone post-translational modification (hPTM) thus marks DSBs and acts as docking site for the recruitment of MDC1 and TP53BP1/53BP1. The latter, together with the XRCC6-XRCC5 proteins and the RIF1-shieldin complex, is responsible for protection of the break sites against end-resection and thereby guides repair pathway choice towards NHEJ. The MRN complex plays a pivotal role as it also facilitates in complex with BRCA1, CtIP and EXO1 extensive end-resection, thus creating single-stranded DNA (ssDNA) filaments rapidly stabilized by the replication factor RPA1. BRCA1 also recruits the PALB2-BRCA2 complex, which loads RAD51 to initiate sister-chromatid strand invasion and DSB repair *via* HR (7).

Because of their central role in cell survival, proteins participating in the DDR are often de-regulated in different types of cancers; interestingly, some loss-of-function mutations represent innovative therapeutic opportunities. The best studied example is BRCA1 deregulation in ovarian and breast cancer, which results in the accumulation of mutations and predisposes to genomic instability (8) while acquiring extreme sensitivity to inhibitors of Poly-ADP-ribose polymerase (PARPi). Indeed, in an HR-deficient background DSB repair largely relies on NHEJ, which requires PARP activity (9). In consequence, HR-defective cells exhibit high sensitivity to PARPi treatment triggering efficient and selective cancer cell death, which is exploited for therapy. Genomic screens are therefore broadly employed to identify synthetic lethality with drugs such as PARPi. Nevertheless, patients often develop drug resistance, thus indicating the need for a deeper characterization of the repair process to rationally propose alternative drug targets.

In light of this, in the past decades the core components of DSB repair pathways have been intensively studied (10-13), however it is only partially understood how these machineries are functionally embedded in the broader chromatin context, and how chromatin determinants impinge on repair pathway choice (14). Beyond γH2AX, the role of other hPTMs in DDR has come into focus (15), either by regulating DNA accessibility and chromatin stiffness (16-19), or by acting as docking sites for the recruitment of DSB repair proteins (20-23). As a consequence, the more accredited models for the DSB repair rely on the coordinated action of determinants belonging to different chromatin layers, including DNA, hPTMs, components of the DDR machineries and chromatin-associated proteins, to harmoniously ensure successful DSB repair. Nevertheless, how these determinants are functionally connected, and how they are dynamically and temporally regulated upon induction of DSBs, are major questions that remain to be resolved. The high complexity of the DDR suggests that no single approach is sufficient to capture the regulation of this fundamental biological process. Therefore, here we bring together the unbiased nature of mass spectrometry with three complementary strategies to study DSB-induced chromatin dynamics at different scales of resolution, namely i) chromatin-wide, by investigating the DNA-bound proteome (iPOC), ii) targeted, by identifying functional interactors of known DDR proteins (ChIP-SICAP) (24, 25), and iii) at the level of mono-nucleosomes, to determine hPTM kinetics (N-ChroP) (26, 27). Moreover, we added a temporal dimension to the study to characterize the DDR process from a chromatin-centered perspective in unprecedented detail and with high temporal resolution. In addition, we validate proteomic data by means of orthogonal functional assays, thus assigning a role of 12 newly identified candidates in the regulation of NHEJ, HR or pathway choice. Moreover, we show that depletion of novel HR-regulating proteins (G9A, ATAD2, TPX2) is synthetic lethal with PARPi. Finally, overlay of these complementary proteomic layers allowed us to reconstruct potential cause-effect mechanisms between DSB-mediated chromatin recruitment and epigenetic regulation during DNA repair. An elective example is represented by the chromatin recruitment of methyltransferase G9A followed by the monomethylation of its substrate H3K56 specifically at *foci*-specific mononucleosomes marked by γH2AX, thus implying a role for this hPTMs in DSB repair. Collectively, beyond providing deep insight into DSB-mediated chromatin dynamics, our approach identifies novel cell vulnerabilities as leads for the development of potential therapeutic interventions. Finally, to facilitate further exploration of chromatin dynamics during the DDR, we make our data available as a resource at www.chromatin-proteomics.dkfz.de.

## RESULTS

### The DSB-mediated interactome dynamics analysis identifies ON-chromatin interactors of MDC1 and RPA upon DSB repair

In this study we applied complementary proteomic approaches to investigate chromatin composition and dynamics during DSB repair, at the level of global chromatin, protein interactions, and hPTMs. First, we employed ChIP-SICAP (24) in cells subjected to ionizing radiation (IR), an efficient inducer of double-strand breaks (DSBs), to identify novel chromatin-bound proteins that are involved in double-strand break (DSB) repair. To this end, we targeted five core components of the DNA repair machinery as bait proteins: the tumor suppressor TP53, the MRN complex subunit RAD50, the HR-associated protein RPA, the mediator of checkpoint MDC1 and 53BP1, important for NHEJ. We used a triple-SILAC labeling approach (28) to distinguish specific binders of target proteins from background in IgG controls, and to accurately quantify DSB-induced changes in the interactomes (Fig.1A). We classified the interactors into either constitutively associated with the bait (constitutive) or exclusively associated while the bait is in its ON-or OFF-chromatin state. Each bait exhibited a different relative proportion of interactors in the three classes; moreover, by performing this analysis in both untreated and cells subjected to IR, we were able to trace the DSB-mediated interactome dynamics. Our data show that, upon DSBs formation, proteins interacting with RPA, MDC1 and 53BP1 have a more dynamic behavior in comparison with binders of p53 and RAD50 that seem to have a more static interactome (Fig.1B). We then evaluated if common chromatin players were recruited at sites marked by the different DDR proteins, or whether each candidate used as bait in ChIP-SICAP had a rather unique interactome. Intersection of the respective binding proteins revealed that roughly 65% of p53, RAD50 and RPA interactors were identified (65%, 69% and 66%, respectively) in association with MDC1. Interestingly, in the same conditions more than 80% of proteins associated with the NHEJ target 53BP1 were co-enriched, while additionally identifying 95 unique binders. Similarly, the vast majority of p53 and RAD50 binders (75% and 86%, respectively) were co-purified in RPA ChIP-SICAP experiments (Fig.1C). Since RPA and MDC1 accounted for more than 90% of the collectively identified interactors, and since RAD50 plays a pivotal role at the junction of the NHEJ and HR repair pathways, we further evaluated the IR-induced changes in the ON-chromatin interactomes of these three targets.

**Figure 1.**
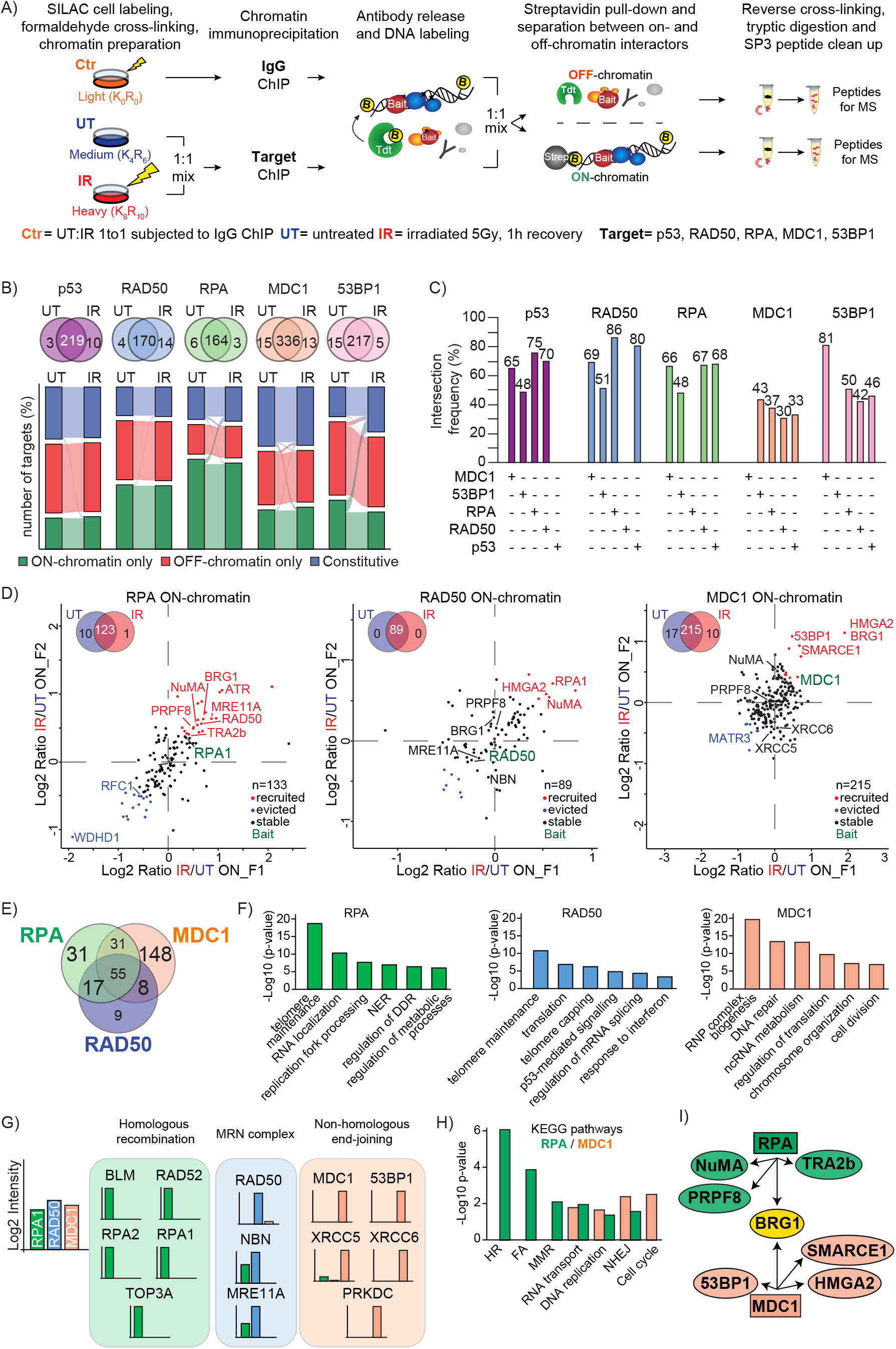
On-chromatin functional interactors of DDR core components. **A**) ChIP-SICAP experimental strategy. Crosslinked chromatin preparation from SILAC labeled cells were enriched for target DDR proteins in the absence (UT) or upon DSB (IR) followed by quantification of functional interactors of the bait while in its ON-or OFF-chromatin state. IgG enrichment serves as internal technical control. **B**) Venn diagrams show the overlap of binders between untreated condition (UT) and upon ionizing radiations (IR). Sankey diagrams represent the IR-mediated dynamics of functional interactors for protein used as bait in A); proteins in common between UT and IR were classified into constitutively associated with the target (Constitutive) or quantified exclusively in the ON- and OFF-chromatin fraction (ON-and OFF-only, respectively). **C**) Bar charts display the frequency of intersection (expressed as percentage) between interactors of the different proteins used as bait in ChIP-SICAP. **D**) Scatterplots representing modulation of ON-chromatin interactors recruited (red) or evicted (blue) from RPA, RAD50, or MDC1 sites upon DSB (IR) in comparison with untreated condition (UT). Venn diagrams show the number of ON-chromatin interactors quantified in each experiment. **E**) Intersection among the ON-chromatin interactors quantified in RPA, RAD50 or MDC1 experiment. **F**) Top-6 gene ontology categories associated with ON-chromatin interactors quantified in RPA, RAD50 or MDC1 ChIP-SICAP. **G**) Log2 intensity of proteins belonging to HR, MRN complex, or NHEJ and enriched in RPA1 (green), RAD50 (blue), and MDC1 (orange) ChIP-SICAP. **H**) KEGG pathways of proteins functionally interacting with RPA (green) or MDC1 (orange), expressed as difference between p-value in RPA and MDC1 (in -log10). **I**) Cartoon representing candidate proteins associated with RPA (green), MDC1 (orange), or both targets (yellow) upon DSB formation. Squares and elliptical shapes correspond to proteins used as bait in ChIP-SICAP, and their DSB-induced interactors, respectively.

In each ChIP-SICAP experiment, the bait was among the most enriched proteins, indicating the specificity of the technique, along with histones, reflecting a successful chromatin enrichment (Extended Data Fig.1A-C). Moreover, between 40 and 75% of the interactors were identified exclusively in the ON-chromatin fraction, thus providing further evidence of the specificity of the approach (Extended Data Fig.1D-F). When using RPA as bait, known components of the HR pathway (e.g. ATR, RPA2 and ETAA1) were strongly recruited on chromatin upon IR, indicating their close proximity to RPA on DNA (Fig.1D). Moreover, specific interactors of RPA belong to gene ontologies (GOs) associated with regulation of DNA damage repair and replication fork processing (Fig.1E, F). In the RAD50 experiment, RPA1 showed strong DDR-induced recruitment, and other proteins that co-enriched with RAD50 are involved in telomere maintenance and p53-mediated signaling (Fig.1D-F). In contrast, ChIP-SICAP for MDC1 identified different core components of the NHEJ machinery (e.g. 53BP1, XRCC5/Ku80 and XRCC6/Ku70), and our results point towards a DNA-mediated interaction among these chromatin proteins (Extended Data Fig.1G-M). Moreover, our data readily revealed also the dynamic ON-chromatin cross-talk between 53BP1 and MDC1, in line with their functional interaction as known readers of γH2AX (Fig.1D, Extended Data Fig.1G). In addition, specific MDC1-interactors enriched GOs terms involved in the DNA repair process (Fig.1D-F). While subunits of the MRN complex were expectedly particularly associated with RAD50, we observed that DNA repair components involved in HR and NHEJ machineries were predominantly enriched in the ON-chromatin fraction of RPA and MDC1, respectively (Fig.1G). The comparison between pathways associated with either RPA-or MDC1-specific ON-chromatin interactors shows how HR and Fanconi Anemia (FA) are extremely enriched in RPA, while interactors of MDC1 belong to cell cycle and NHEJ (Fig.1H). These results are in line with the notion of MDC1 as key regulator of the DDR cascade, in contrast with the HR-restricted role of RPA.

Besides known components of the DDR machinery, our analysis identified numerous other proteins interacting with MDC1 or RPA upon IR, but not previously associated with a function in the DDR. Among these, we focused on several candidates to functionally assess their role in DSB repair through either HR or NHEJ repair pathways. From our ChIP-SICAP data, we selected NuMA, TRA2B and PRPF8 as HR-related candidates specifically associating with RPA upon DSB formation, and SMARCE1/BAF57 and HMGA2 among the proteins co-enriched with MDC1. Moreover, we investigated the role of SMARCA4/BRG1 in light of its strong functional DDR-mediated interaction with both RPA and MDC1 (Fig.1I).

### ON-chromatin interactors of RPA and MDC1 influence DNA repair by regulating HR, NHEJ or DSB repair pathway choice

To assess the involvement of IR-dependent functional interactors of RPA and MDC1 in DSB repair, we first evaluated whether their silencing had any impact on the formation and resolution of γH2AX *foci*, an early marker of DSB repair. To this end we took advantage of isogenic U2OS AID-DIvA cells, where the AsiSI enzyme produces a defined number of DSBs upon 4-OHT induction, while the DSB repair is promoted through Auxin-Inducible degradation of the enzyme (7). Upon target knockdown (Extended Data Fig.2A), all tested cells show a substantial decrease in the number of induced γH2AX *foci* and, besides silencing of the positive control RPA and 53BP1, only depletion of HMGA2 still preserves a statistically significant DSB response (Fig.2A). These results therefore suggest that all tested candidates identified *via* ChIP-SICAP are crucial for either γH2AX *foci* formation or spreading of this marker (e.g. HMGA2) upon IR, implicating their function in DSB repair.

To gain insight into the role of these candidates in DSB repair by HR or NHEJ, we employed isogenic U2OS-traffic-light reporter (U2OS-TLR) cells (29), and used FACS-based quantification to simultaneously evaluate which of the two main DSB repair pathways is impaired upon target knockdown. Cells depleted for 53BP1 and RPA/RAD51 were used as positive controls for NHEJ and HR, respectively (Fig 2B). Silencing of HMGA2 results in a strong increase in the number of cells repairing DSBs *via* HR, while the fraction of NHEJ remains unchanged (Fig.2B, Extended Data Fig.2B, C). In contrast, silencing of BRG1 specifically decreases the efficiency of HR, while inhibition of TRA2B promotes the rate of NHEJ by 20% (Fig.2B, Extended Data Fig.2B, C). These observations clarify the controversial role of HMGA2 in the DDR (30) and indicate how this target binds MDC1 and has a similar function to 53BP1 in negatively regulating HR. In line with previous evidence (31) our results also suggest that BRG1 is recruited at DSBs to promote a chromatin state facilitating HR repair (Fig.2B, Extended Data Fig.2B, C). In addition, besides its canonical function in BRCA1 RNA splicing (32), our results suggest a novel and more direct role for TRA2B in the DDR *via* inhibition of NHEJ.

**Figure 2.**
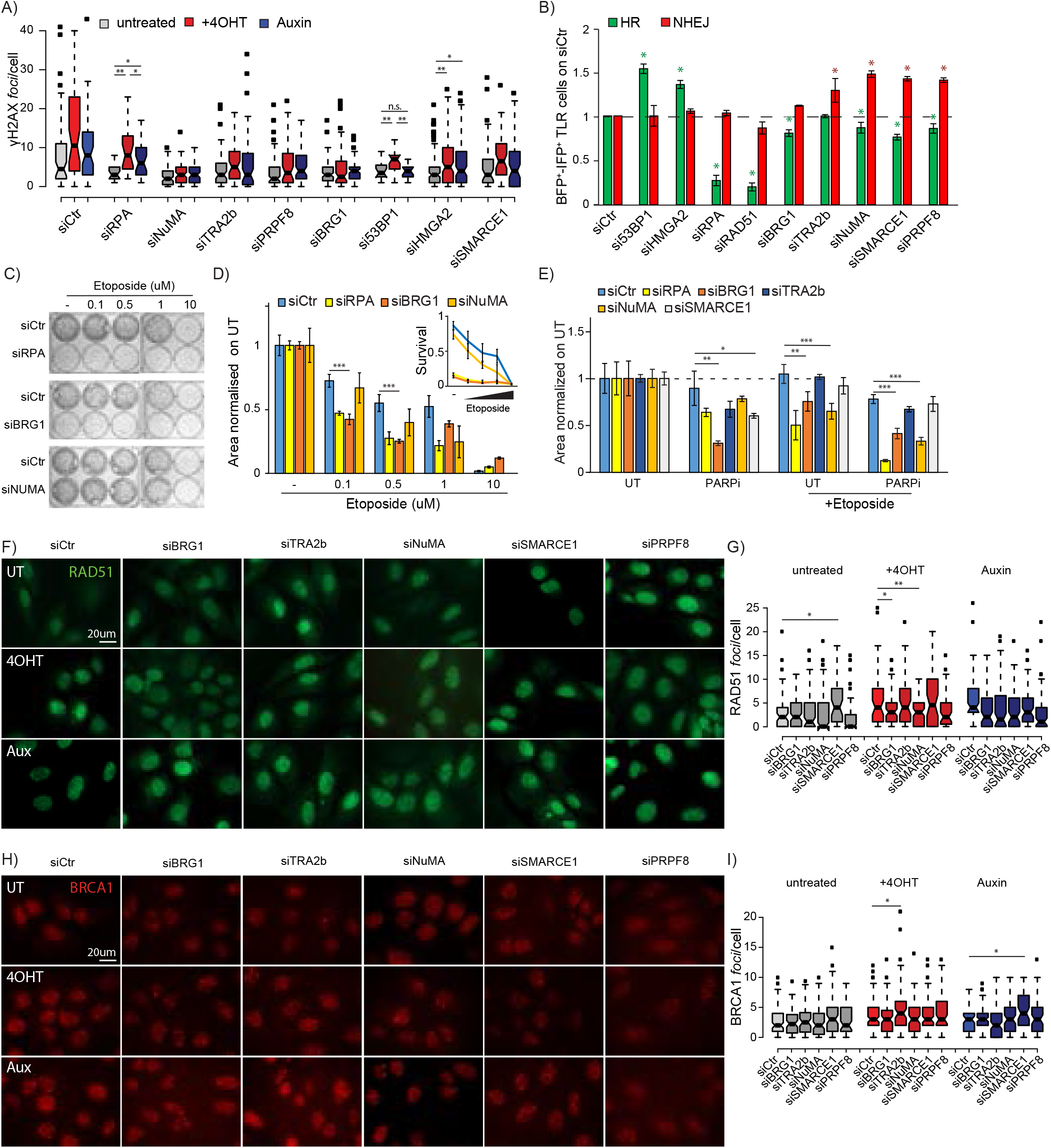
Functional characterization of RPA-and MDC1-specific on-chromatin interactors. **A**) Number of γH2AX *foci* per nucleus in AID-DIvA cells left untreated, upon DSB induction, and during DSBs repair (untreated, +4OHT and Auxin, respectively) upon knockdown for targets interacting with either RPA or MDC1 in comparison with on-target non-targeting silencing control (siCtr). **B**) Quantification of homologous recombination (green) and non-homologous end joining (red) repair events in traffic-light reporter (U2OS-TLR) cells depleted of protein candidates and normalized on silencing control (siCtr). Green and red asterisks reflect significant regulation in HR and NHEJ, respectively, in respect to siCtr (ANOVA-based statistics). Mean values of at least three biological replicates with standard deviation are shown. **C**) Colony formation assay at increasing amounts of etoposide in cells knockdown for proteins important for HR repair (BRG1) or pathway choice (NuMA) compared with silencing control. Silencing of RPA (siRPA) is used as internal control. **D**) Quantification of C, mean values normalized on untreated condition (UT) and standard deviations are reported. Zoom-in shows the survival of cells upon target knockdown compared with silencing control. Mean values of at least three biological replicates with standard deviation are shown. **E**) Quantification of colony formation assay in U2OS cells depleted of listed proteins involved in either HR or pathway choice and subjected to PARP (PARPi) alone or in combination with etoposide to promote DSBs formation. Mean values from three biological replicates normalized on untreated condition (UT) are shown, error bars represent standard deviations. Representative immunofluorescence images of RAD51 (**F**) and BRCA1 (**H**) *foci* in AID-DIvA cells in untreated (UT) condition, upon DSB formation (4OHT), and during DSB repair (Aux) upon knockdown with indicated targets. siCtr corresponds to on-target non-targeting siRNA. Quantification of RAD51 (**G**) and BRCA1 (**I**) *foci* in AID-DIvA cells knockdown for targets involved in either HR or pathway choice. *, ** and*** correspond to p-value < 0.05, 0.01 and 0.001 of ANOVA statistical test, respectively. n.s. = not statistically significant.

In contrast to most site-specific DSB repair systems, the U2OS-TLR assay allows to simultaneously quantify how individual proteins impact on the equilibrium between NHEJ and HR at induced DSBs. In particular, knockdown of NuMA, SMARCE1 or PRPF8 leads to an increase in NHEJ mirrored by a parallel decrease in HR (Fig.2B, Extended Data Fig.2B, C). These results add functional detail to the suggested involvement of NuMA and PRPF8 in DNA damage repair (35-37), while assigning a completely novel role in the DDR to the BAF/PBAF subunit SMARCE1.

We next selected BRG1 and NuMA as representative proteins important for HR and repair pathway choice, respectively, and evaluated whether their involvement in either processes can be effectively distinguished. Knockdown of BRG1, but not NuMA, significantly sensitized cells to low doses of the DSB-inducer etoposide (from 0.1 to 0.5µM), to a similar extent as RPA depletion which was used as a control for HR impairment (Fig.2C, D). This corroborates the results obtained in the TLR assay (Fig.2B) and indicates how, upon depletion of a pathway choice repair protein (e.g. NuMA), the increased rate of NHEJ confers partial resistance to low doses of DSB inducer.

Selective impairment in the HR pathway is currently exploited in cancer therapy due to the predisposition to synthetic lethality with PARP inhibitors (PARPi). We therefore challenged our results obtained from the TLR system by exposing cells knockdown for the different targets to Olaparib treatment, either alone or in combination with etoposide. As expected, silencing of HR-related proteins (i.e. BRG1, RPA) resulted in pronounced synthetic lethality with PARPi, while knockdown of targets involved in repair pathway choice only partially sensitizes to either Olaparib (i.e. siSMARCE1) or etoposide treatment (i.e. siNuMA) (Fig.2E). These results strongly suggest that, in contrast to HR-deficient cells, the increase in NHEJ observed upon knockdown of a DSB pathway choice regulator might counterbalance the deficiency in HR, thereby decreasing the sensitivity to PARPi.

To gain further insights in the function of selected target proteins in DSB repair, we monitored the formation of RAD51 and BRCA1 *foci*, in AID-DIvA cells. Our results revealed that NuMA-depleted cells are defective in RAD51 loading, while siTRA2B cells have a significantly higher number of induced BRCA1 *foci*, thus providing a potential explanation for their different role in DSBs repair (Fig.2F–I, Extended Data Fig.2F, G). Importantly, none of these results can be explained by a change in cell cycle distribution upon depletion of the target proteins (Extended Data Fig.2D, E).

In summary, we investigated the potential role in DSB repair for candidates associating with either RPA or MDC1 and highlighted that inhibition of each tested candidate has an impact on the DDR as monitored by γH2AX *foci* formation. Moreover, we assigned a role to these targets in NHEJ (i.e. HMGA2), HR (i.e. BRG1, TRA2B), or the balance between these DSB repair pathways (i.e. NuMA, SMARCE1, PRPF8). As validation, knockdown of HR-associated targets, but not of proteins involved in the repair pathway choice, shows synthetic lethality with PARPi. Finally, we show that although NuMA and TRA2B are both enriched at RPA sites in ChIP-SICAP, they exhibit functional differences, thereby suggesting that mediators of the DDR can impinge on DNA repair in distinct ways.

### Dynamic profiling of chromatin-associated interactors of γH2AX

Gamma-H2AX is the first marker of DSB repair upstream of both NHEJ and HR pathways (39), and MDC1 has been identified as its main reader (38). However, little is known about the identity and dynamics of other chromatin proteins that associate with this fundamental hPTM. To investigate this in a temporally resolved manner, we applied ChIP-SICAP to the hPTM γH2AX, thereby using this approach for a PTM for the first time. We profiled the dynamics of its ON-chromatin interactors in a time course during the DSB repair induced by IR, and employed triple-SILAC labeling to discriminate, at each time point, γH2AX-specific ON-chromatin interactors from general histone (H2A)-associated proteins and from non-specific background (IgG isotype control) (Fig.3A). Quantitative analysis identified almost 150 interactors that exclusively associate with γH2AX, including the strong and IR-dependent enrichment of the γH2AX-reader MDC1, illustrating the efficacy of the time-course ChIP-SICAP approach, also when applied to hPTMs (Fig.3B, C, Extended Data Fig.3A-E).

Among the γH2AX-specific proteins, almost half are linked to regulation of different chromatin processes, and 25 have an already established role in DNA repair, in addition to proteins with a canonical role in various other biological processes (Extended Data Fig.3F, G). Furthermore, γH2AX-specific interactors identified at all time-points along the repair process are enriched in SUMO E3 ligases (Extended Data Fig.4A), in line with the well-established role of SUMOylation in DNA repair (40). In the same group, we identified proteins involved in RNA splicing (Extended Data Fig.3H), thus suggesting a possible stabilization of DSB-induced transcripts (41). In contrast, proteins evicted from γH2AX-marked sites were more globally involved in ribosome formation and regulation of translation (Extended Data Fig.3I).

Quantitative analysis of ChIP-SICAP data revealed that almost 1/3 of γH2AX interactors are shared with its reader MDC1 (98 proteins), of which 72 are exclusively identified by these two chromatin determinants, and that only a minor fraction is in common with the other DSB-related proteins (Fig.3D). Moreover, our data allow to profile the temporal association between γH2AX and ON-chromatin interactors identified in MDC1, RPA and RAD50 ChIP-SICAP experiments, thus both validating the chromatin association for these binders at damaged sites, and describing their dynamics during the DDR at sites marked by γH2AX (Fig.3E – G). Overall, the majority of proteins are rapidly recruited at DSB sites, although they follow different kinetics during the DDR. In particular, interactors of MDC1 show the highest dynamics at break sites with a rapid chromatin recruitment, in line with the strong link between γH2AX and its reader (Fig.3E). Most of the RPA and RAD50 binders exhibit a lower temporal enrichment at γH2AX-sites, and in general they are recruited more gradually during the repair process (Fig.3F, G).

Across the proteins that associate with γH2AX, our data indicate how proteins with a similar function can follow a different kinetics at break sites, and, conversely how targets belonging to distinct biological processes can associate with γH2AX at the same time during the DDR. This emphasizes the complex regulation of chromatin around the break site to ensure an optimal repair process (Fig.3H). Beyond the many ON-chromatin interactors of γH2AX with an already established link to the DDR (purple in Fig.3H), we identified several others that are less thoroughly investigated in this context. This includes the nuclear protein THRAP3, the histone reader ATAD2 and the microtubule organizer TPX2, whose role in the repair process was further investigated in more detail.

In summary, our time course ChIP-SICAP experiment performed during the DDR and using γH2AX as entry point revealed a functionally and temporally diverse set of proteins, indicating intricate chromatin re-arrangements at DSB sites. Among many proteins previously associated with DDR, this approach also identified novel candidates whose function in DNA repair remains to be determined.

### THRAP3 is rapidly recruited to γH2AX sites upon DNA damage

THRAP3 caught our interest because of its recruitment to γH2AX-marked break sites predominantly at 1h after inducing DSB (Fig.3C, Extended Data Fig.3A–E). THRAP3 was previously reported in an overexpression system to be highly phosphorylated upon DNA repair but to be excluded from micro-irradiated lesions (42). More recently it has been implicated in R-loop resolution through the interaction with DDX5 and XRN2 (43). Our analysis showed that all these three factors associate specifically with γH2AX, albeit at different time points during the DDR (Supplementary Information 1), suggesting an additional role of THRAP3 beyond R-loop resolution. We therefore first tested the co-localization of endogenous THRAP3 with γH2AX in untreated as well as irradiated cells by means of immunofluorescence (IF) and proximity ligation assay (PLA). Our results verified the interaction between these two chromatin determinants already in untreated conditions, and confirmed the significant increase in co-localization between THRAP3 and γH2AX after 1h recovery upon induction of DSBs (Fig.4A). Furthermore, silencing of THRAP3 impaired γH2AX *foci* formation but not their repair in AID-DIvA cells system (Fig.4D, Extended Data Fig.4A, C). Moreover, TLR-U2OS cells show a normal balance between NHEJ and HR after knock-down of THRAP3 (Fig.4E, Extended Data Fig.4E). Taken together, our results suggest that THRAP3 is specifically but transiently associated with γH2AX. Functionally, it affects both DSB repair pathways leaving the balance between NHEJ and HR untouched. These results, point towards a more structural role and possible involvement of THRAP3 in either promoting the splicing of mRNA encoding for repair proteins important for γH2AX *foci* formation, or in the stabilization of DNA damage-induced RNAs at DSB sites.

### ATAD2 and TPX2 stably interact with γH2AX during DNA damage repair and play a key role in HR

Both the specific γH2AX-reader MDC1, as well as the topoisomerase II alpha and the helicase DDX21, two enzymes involved in DNA and R-loop unwinding and regulating genome stability (44, 45), were strongly enriched at γH2AX-marked DSBs at all time-points during the DDR (Fig.3C, E, H; Fig. 4A). This encouraged us to further investigate proteins with a similar temporal trend at γH2AX sites, yet with a poorly characterized role in DDR. In particular, within this cluster of proteins we focused our attention on the spindle protein TPX2 and the transcriptional co-activator ATAD2 (Fig.3B, C, H, 4A, Extended Data Fig.3A–E). Upon IR, TPX2 has been reported to accumulate at DSBs to negatively regulate 53BP1 (46, 47), however little is known about its role in the DDR. ATAD2 binds in vitro to hyper-acetylated histone H4 tails through its bromodomain (48), and it is transcriptionally induced by anti-cancer and DNA-damaging agents *via* ATM and ATR checkpoint kinases. Silencing of ATAD2 sensitizes triple-negative breast cancer cells to carboplatin treatment (49, 50) but its role in DSB repair has been not investigated before.

**Figure 3.**
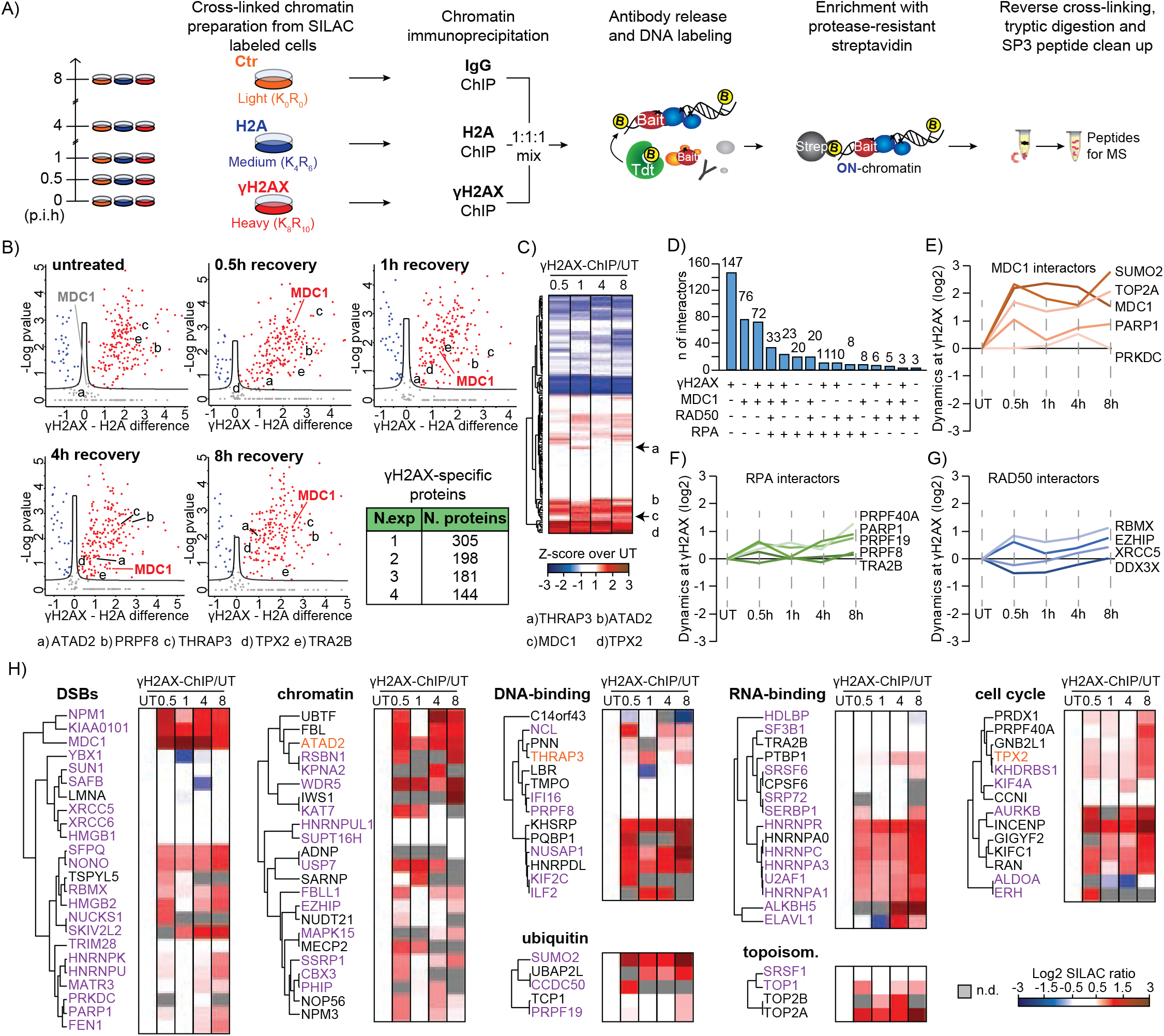
Functional interactors of γH2AX during DNA damage repair. **A**) Time course ChIP-SICAP experimental design; p.i.h.: hours post-irradiation. At each time point U2OS cells labeled with light-, medium-or heavy-SILAC amino acids were subjected to ChIP-SICAP protocol using IgG control, H2A or γH2AX as bait, respectively. **B**) Volcano plots represent the fold change difference of t-test statistics for γH2AX- and H2A-associated ON-chromatin binders (red and blue, respectively), at different time points during DSBs repair. MDC1 is highlighted as positive control. The table represents the number of γH2AX-specific functional interactors shared among the different experiments during DSBs repair (from untreated condition to 8h). a to e refer to listed γH2AX-specific ON-chromatin functional candidates. **C**) Heatmap representation of γH2AX-specific ON-chromatin interactors identified at all time points during the DDR. At each time point, log2 ratios over untreated sample are shown upon z-score normalization. Red and blue correspond to enriched and evicted γH2AX interactors during DSBs repair, respectively. a to d refer to listed ON-chromatin candidates further evaluated. **D**) Overlay view of number of interactors in common among γH2AX and the DDR-associated proteins used as bait in ChIP-SICAP in figure 1. Line plots represent the dynamics of MDC1-(**E**), RPA-(**F**), or RAD50-specific (**G**) ON-chromatin functional interactors at γH2AX-sites during the DDR. **H**) Heatmap visualization of γH2AX-specific ON-chromatin interactors classified according to their function; SILAC ratio (in log2) between the protein abundance at different time points during the DDR and untreated conditions is shown. Proteins already associated with DSB break repair and novel candidates for γH2AX are shown in purple and orange, respectively. n.d. = not determined.

**Figure 4.**
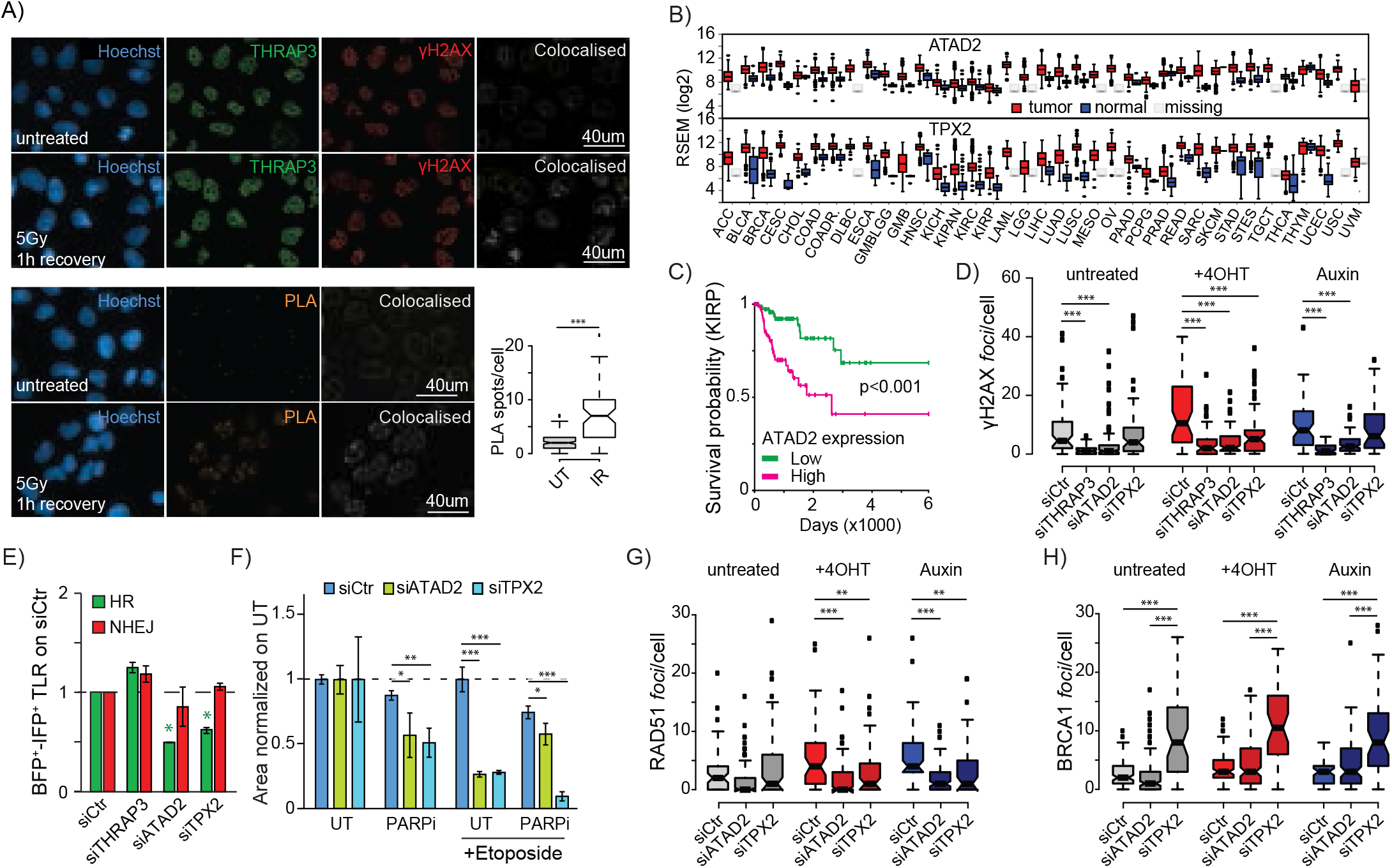
IR elicits functional interaction between THRAP3 and γH2AX, while ATAD2 and TPX2 mediate DSB repair through HR. **A**) Immmunoflurescence (top panel) and proximity ligation assay (PLA, bottom panel) validation of DSB-mediated interaction between THRAP3 and γH2AX. Boxplot shows the quantification of PLA assay in untreated cells (UT) and cells exposed to ionizing radiation (IR, 5Gy, 1h recovery). **B**) Deregulation of ATAD2 (top) or TPX2 (bottom) RNA level in tumor samples (red) compared to normal tissue (blue) from TCGA expressed as log2 RNAseq by Expectation-Maximization (or RSEM). **C**) Survival probability of patients affected by Kidney renal papillary cell carcinoma (KIRP) with low (green) or high (pink) ATAD2 levels. **D**) Number of γH2AX *foci* per nucleus in AID-DIvA cells left untreated or subjected to DSBs induction and repair (tamoxifen/+4OHT and Auxin, respectively), after knockdown for THRAP3, ATAD2 or TPX2 in comparison with on-target non-targeting silencing control (siCtr). Significance over siCtr is shown. **E**) Quantification of HR (green) and NHEJ (red) repair events in U2OS-TLR cells depleted for THRAP3, ATAD2, or TPX2 and normalized over non-targeting silencing control (siCtr). Green and red asterisks reflect significant regulation in HR and NHEJ, respectively, in respect to siCtr. **F**) Quantification of CFA in U2OS cells depleted of either ATAD2 (light green) or TPX2 (light blue) and subjected to PARP inhibitor (PARPi) alone or in combination with etoposide to promote DSBs formation. Mean values of three replicates normalized on untreated (UT) with standard deviations are shown. Quantification of RAD51 (**G**) and BRCA1 (**H**) *foci* in AID-DIvA cells knockdown for targets involved in either HR or pathway choice. *, ** and*** correspond to p-value < 0.05, 0.01 and 0.001 of ANOVA statistical test, respectively.

As derived from the TCGA database, both TPX2 and ATAD2 are overexpressed or amplified in aggressive tumors (Fig.4B), and high levels of ATAD2 correlate with poor prognosis especially in breast and kidney cancer (Fig.4C). These observations, together with the fast chromatin recruitment of both ATAD2 and TPX2 at γH2AX sites upon DSB induction, prompted us to further elucidate their role in DSB repair. Knockdown of either target resulted in a significant impairment in γH2AX *foci* formation and repair (Fig.4D, Extended Data Fig.4B, C). In particular, while ATAD2-deficient cells failed to mount a correct early DSB response, depletion of TPX2 resulted in a progressive accumulation of unrepaired DSBs (Fig.4D, Extended Data Fig.4C). To further address their role in DSB repair, we adopted the U2OS-TLR cell system and identified a dramatic impairment of over 50% in HR efficiency upon silencing of either ATAD2 or TPX2 (Fig.4E, Extended Data Fig.4D, E). Notably, the severity of the effect almost reaches the level observed upon depletion of the hallmark HR repair proteins RPA and RAD51 (Fig 2B). We therefore tested whether knockdown of ATAD2 or TPX2 could confer synthetic lethality in a combined treatment with PARPi (Fig.4F). Our results show that cells deficient for either of the targets were sensitive to PARPi, thus validating their role in HR-mediated DSB repair. Interestingly, this effect was synergistic with etoposide treatment in TPX2-but not in ATAD2-depleted cells (Fig.4F), thereby implying that these two candidates might play a different role in the HR cascade. Indeed, while knockdown of either ATAD2 or TPX2 affected the formation of induced RAD51 *foci* (Fig.4G, Extended Data Fig.4H), in line with the reported interaction with the cell cycle regulator Aurora kinase A (AURKA) (47), only TPX2 silencing resulted in a pronounced accumulation of cells in G2/M phase and a significant increase of BRCA1 *foci* (Fig.4H, Extended Data Fig.4F, G, I).

Collectively, we identified ATAD2 and TPX2 as novel interactors of γH2AX, and provide evidence for key roles for both proteins in HR-mediated repair. Moreover, both ATAD2 and TPX2 might represent promising drug targets conferring synthetic lethality with PARPi in HR-proficient cells.

### Characterization of DNA repair-induced chromatin dynamics through Isolation of Protein On Chromatin (iPOC)

ChIP-SICAP experiments represent a candidate approach, requiring *a priori* knowledge for the target protein used as a bait to enrich associated chromatin proteins. To complement the view that we obtained in this way for γH2AX, RPA, RAD50 and MDC1, we therefore aimed at extending this vision by developing an innovative and unbiased approach to determine DSB-induced changes in overall chromatin protein composition. Similar to recent strategies developed to study proteins interacting with nucleic acids (51-56), we exploited the use of nucleotide mimetics as a tool to mark the DNA, leaving a chemical trace amenable for the selective isolation of <u>p</u>rotein <u>o</u>n <u>c</u>hromatin (or iPOC). In particular, we labeled the DNA *via* full incorporation of 5-Ethynyl-2’-deoxyuridine (EdU) for subsequent capture *via* copper-catalyzed azide-alkyne cycloaddition (CuAAC) of biotin azide (Fig.5A, Extended Data Fig.5A, B). We combined this methodology with triple SILAC protein labeling and mass spectrometry to precisely quantify changes in chromatin composition during the DDR. Specifically, medium-or heavy-labeled cells were subjected to EdU labeling and then either left untreated or collected at different time points (i.e. 1h, 4h, 8h) after DSB induction by IR, respectively. Light-SILAC cells were not subjected to EdU labeling and served as negative control. At each time point, crosslinked cells from the three differently SILAC-labeled samples were mixed in equal amounts and subjected to click chemistry-based biotin binding; chromatin was then sheared and DNA-binding proteins were enriched by means of protease-resistant streptavidin beads (or prS) (25). Upon tryptic digestion and mass spectrometry analysis, we characterized the dynamics of chromatin-binding proteins during DSB repair (Fig.5B). To our knowledge, it is the first time that a similar approach has been employed in a quantitative and time-course manner to describe global changes in chromatin composition.

Protein quantification showed the efficient and highly reproducible enrichment (≥ 0.85) of DNA-binding proteins identified in EdU-treated samples over the negative control (Extended Data Fig.5C, D). Moreover, the high Spearman correlation (> 0.6) between protein abundances at different time points indicates that iPOC is a very sensitive strategy that enables capturing the highly dynamic nature of chromatin during the DSB response (Extended Data Fig.5E). To further asses the sensitivity, we benchmarked iPOC against deeply fractionated chromatin input where almost 4000 proteins were profiled during the DSB response (Extended Data Fig.5F, G). iPOC quantified almost 200 proteins that were either not identified in any of the ChIP-SICAP experiments or that were exclusively enriched in iPOC (i.e. 123 and 66 proteins, respectively) (Fig.5C). These results highlight the orthogonality between the investigation of DNA-binding determinants associated with a target protein involved in DSB repair, and the unbiased dissection of chromatin regulation during DNA repair. Interestingly, iPOC-specific proteins were mainly enriched in gene ontology terms involved in chromatin organization, DSB repair and chromatin post-translational modification, thus further stressing the functional role of determinants exclusively quantified by iPOC (Fig.5D). In line with this, different classes of proteins associated with histone, transcription, chromatin regulation, DNA damage repair, and post translational modification (i.e. ubiquitin) were over-represented in iPOC in comparison with the chromatin input (Fig.5E), thereby indicating the high specificity of iPOC in enriching DNA-binding proteins. Accordingly, while proteins deregulated during the DSB response in the chromatin input were mainly involved in RNA processing and cell cycle (Extended Data Fig.5H, I), candidates quantified in iPOC at 1h, 4h or 8h time points were enriched in GO terms associated with DNA repair and chromatin organization (Fig.5F). Moreover, the vast majority of proteins identified in iPOC were either not detected or not significantly deregulated in the chromatin input (Fig.5G, Extended Data Fig.5J), indicating that enrichment of chromatin-bound proteins *via* iPOC enhances sensitivity in detecting the dynamics of chromatin-associated proteins.

In iPOC we identified the DSB-induced chromatin recruitment of different classes of epigenetic regulators ranging from histone modifying enzymes, to structural and core components of molecular machineries regulating the DDR. Moreover, in contrast to the chromatin input, in iPOC we observed the chromatin enrichment for candidates identified *via* ChIP-SICAP experiments such as MDC1, 53BP1, BRG1, NuMA, ATAD2 and TPX2. This result further corroborates the DSB-mediated chromatin association of novel candidates, while highlighting the sensitivity and specificity of our approach in detecting expected known DSB markers. For example, iPOC identified DNA repair proteins that associate with chromatin upon IR such as the topoisomerase II B (TOP2B), the telomeric repeat-binding factor TERF2 and the NHEJ regulator RIF1. Interestingly, upon DSB induction, we observed an overrepresentation of chromatin-modifying enzymes among iPOC-enriched proteins, including subunits of acetyltransferase complexes (e.g. MORF4L1, KAT7 and BRD1), the histone demethylase KDM2A, the hPTM reader and transcriptional regulator PSIP/LEDGF, and the methyltransferases KMT2B/MLL4, WHSC1/NSD2, SETMAR and EHMT2/G9A (Fig.5G, Extended Data Fig.5J). Taken together, these results demonstrate the sensitivity of iPOC in capturing the DNA damage-induced dynamics even for proteins with very fast kinetics such as chromatin remodelers.

Moreover, and in contrast to targeted approaches like ChIP-SICAP, iPOC has the inherent ability to investigate the overall histone composition in chromatin. We observed that in the chromatin input the abundance of most histones declined upon IR, while this effect was generally less pronounced for DNA-bound histones identified through iPOC (Extended Data Fig.5K). Our data are therefore in line with the recently reported proteasome-mediated depletion of histones during DNA repair (31), but suggest that soluble histones might strongly contribute to this phenomenon. Even more interestingly, in iPOC we observed that changes in DNA-bound histones during DSB repair occurred in a time-and histone variant-specific manner, sometimes even leading to a transient increase (e.g. H1.4, H2A.1-3) (Extended Data Fig.5K). These results therefore point towards a mechanism to retain, evict or recruit distinct variants to potentially drive a chromatin conformation status amenable for the DSB repair process, mirrored by the dynamic recruitment of histone chaperones observed in iPOC (Fig5G).

Collectively, the iPOC approach allows for the unbiased temporal characterization of the chromatin composition and the interplay among its regulators during DNA repair, thus providing an orthogonal view point to the candidate-approaches based on ChIP-SICAP. In addition, our data report a strong overrepresentation of chromatin modifying enzymes and proteins involved in post-translational modifications of histone and non-histone proteins, thereby highlighting the sensitivity of the methodology in capturing the dynamic enrichment of transient chromatin interactors.

### ADNP, SMARCA1 and PHF14 are novel chromatin-associated proteins with distinct functions in DSB repair

Among the candidates identified as recruited at chromatin exclusively *via* iPOC, we selected three targets belonging to different functional groups to assess their potential role in DNA damage repair (Fig.5G). In particular, we focused on two negative prognostic markers in cancer, ANDP and PHF14 (57, 58), and a chromatin regulator, SMARCA1/SNF2L (59). We first evaluated the formation and repair efficiency of γH2AX *foci* in cells after knockdown of these targets (Extended Data Fig.5L). We observed that silencing of ADNP increased the number of *foci* already in untreated conditions, resulting in the accumulation of unrepaired DSBs (Fig.5H, Extended Data Fig.5M). Recent evidence showed that ADNP, together with CHD4 and HP1, forms the ChAHP remodeling complex involved in the regulation of higher-order chromatin structure (60). The DSB-induced recruitment of CHD4 observed in iPOC might therefore point towards a possible role of this complex in genome stability, but we cannot exclude a separate function of ADNP during DDR. In agreement with our observation on the role of ADNP in DSB repair, depletion of ADNP in U2OS-TLR cells affected NHEJ by negatively regulating DSB repair *via* HR (Fig.5I, Extended Data Fig.5N, O), an effect that was not caused by cell cycle deregulation (Extended Data Fig.5P, Q). In contrast, similar experiments demonstrated that SMARCA1 and PHF14 are both involved in homologous recombination pathway by preventing NHEJ repair (Fig.5I, Extended Data Fig.5N, O). We therefore suggest that upon depletion of these two candidates, the higher rate of the fast end-joining repair might explain the lower number of unrepaired γH2AX *foci* (i.e. Auxin-treated sample) in comparison with silencing control (Fig.5H, Extended Data Fig.5M). Accordingly, silencing of neither SMARCA1 nor PHF14 promoted synthetic lethality with PARPi, as observed upon depletion of TRA2B (Fig.2E), which is similarly involved in inhibition of NHEJ.

Interestingly, although having a similar role in DNA repair, PHF14 and SMARCA1 seem to follow a different kinetics at chromatin, suggesting a potential non-redundant function. Indeed, depletion of PHF14 sensitized cells to the chemotherapeutic agent etoposide and caused defective formation of RAD51 *foci* (Fig.5J, K), corroborating recent findings (61). In contrast, we did not observe a significant impact of SMARCA1 depletion on either RAD51 or BRCA *foci* (Fig.5K, L, Extended Data Fig.5R, S), thus suggesting a more structural role for the core component of the Nucleosome remodeling factor (NuRF) during DSB repair.

In summary, iPOC identified multiple proteins that associate with chromatin upon DSB, and the functional characterization of selected candidates (e.g. SMARCA1, PHF14 and ADNP) provide evidence for their involvement and mechanism in the DDR, thus underscoring the discovery power of this methodology.

### Dynamics of hPTMs in chromatin and at DSB-sites during DNA damage repair

The overrepresentation of chromatin-modifying enzymes, in particular methyltransferase observed in iPOC (Fig.5G), together with their established fundamental function in DNA damage repair (62), prompted us to investigate histone PTMs as an additional regulatory layer that impinges on chromatin stiffness and accessibility. In particular, we employed Native Chromatin Proteomics (N-ChroP) (26, 27) in a time-course manner to globally characterize the dynamic changes of hPTMs during the DDR (hereafter termed DDR-induced hPTMs) with single nucleosome resolution. In parallel, we profiled temporal trends of histone modifications specifically at γH2AX-containing mononucleosomes (henceforth *foci*-specific hPTMs).

To achieve this goal, we prepared mononucleosomes from U2OS cells at different time points during IR-induced DSB repair and we profiled the hPTM-dynamics *via* the ArgC-like in-gel digestion prior to mass spectrometry analysis (Extended Data Fig.6A, B). For each hPTM, we described its relative abundance (RA) compared with the amount of concomitant modifications on the same peptide, and the relative enrichment (RE) corresponding to the fold enrichment between the RA of a certain modification in γH2AX-mononucleosomes and in the native chromatin mononucleosomes preparation (i.e. input) (Fig.6A). As a result, we identified 33 different histone modifications on 14 peptides from histone H2A, H3 and H4, and we quantified their temporal abundance during DSB repair (Fig.6B, Extended Data Fig.6C– E). To our knowledge, this represents the first example of extensive and unbiased temporal profiling of hPTMs during the DDR with single-nucleosome resolution.

Our analyses indicate that already in untreated conditions γH2AX-mononucleosomes have a distinct hPTM pattern, being enriched for hyper-acetylated in the N-terminal tail of histone H4, a modification associated with open chromatin state (63), and mono-methylated lysine 20 on histone H4 (H4K20me1) (Fig.6B), a known hPTM with a pivotal role in DSB repair (64, 65). These results suggest that γH2AX-containing nucleosomes may be inherently enriched in modifications that establish a more relaxed chromatin state, to potentially promote accessibility for the repair machinery and thus “prime” for DNA damage repair signaling.

In addition, our analysis allows distinguishing hPTMs that are globally induced upon DSB (DDR-induced), from modifications acquired exclusively in mononucleosomes in close proximity to the break site and marked by γH2AX (*foci*-specific). Among the DDR-induced hPTMs, we identified that dimethylated K79 on histone H3 (H3K79me2) has a bimodal enrichment at 30min and 4h (Fig.6B, Extended Data Fig.6D). The abundance of this modification positively correlates with transcriptional rate (66) and can supposedly act as docking site for 53BP1 (67). The synchronous DSB-mediated increase of this modification in both native chromatin input and in γH2AX-mononucleosomes, suggests that H3K79me2 might globally promote the transcription of DNA repair genes, while at DSB sites it could stimulate the production of DDR-induced RNAs. In the same class of modification, we interestingly identified that K95 of histone H2A (H2AK95) is globally monomethylated during DNA damage repair and accumulates at late time points (Fig.6B, Extended Data Fig.6C). Although little is known about modifications on this core histone residue (68, 69), our results suggest involvement of H2AK95me1 in the DDR signaling.

Recent efforts have been dedicated to the investigation of *foci*-specific hPTMs, the most successful advance relies on the intersection between breaks-labeling and sequencing (BLESS) profiles and ChIP-sequencing tracks of *a priori* selected hPTMs (15, 70). While BLESS distinguished hPTMs enriched at NHEJ-and HR-prone DSBs, this approach falls short in exploring unpredicted modifications and in studying their combinatorial occurrence. Our proteomic-based approach circumvents this by the quantitative, unbiased and simultaneous analysis of multiple hPTMs. Indeed, at γH2AX-mononuclesomes we observed an increase in mono- and di-methylation on K20 of histone H4 (H4K20me1, H4K20me2) (Fig.6B, Extended Data Fig.6E), two modifications with a pivotal role in DSB repair pathway choice (21-23). In particular, our time-course analysis showed different kinetics between mono- and di-methylation, where H4K20me1 precedes the dimethylation, enriched at later time points. This effect could either be due to the sequential steps required for achieving a higher methylation degree, or potentially to a different role of H4K20me1 and H4K20me2 in the DDR cascade.

Encouraged by the characterization of dynamics for expected hPTMs at break sites, we expanded our analysis to unexpected trends during the DDR. Among the *foci*-specific hPTMs we observed a progressive increase of monomethylation of K18 and acetylation of K64 on histone H3 (H3K18me1 and H3K64Ac, respectively) (Fig.6B, Extended Data Fig.6D). Monomethylation of H3K18 is mutually exclusive with the acetylation on the same residue and it has been associated with silencing chromatin (72). In DNA repair, H3K18 needs to be deacetylated through SIRT7 to allow 53BP1 recruitment at DSBs, therefore our results suggest that H3K18me1 is a rapid and possibly important mediator of NHEJ repair. In contrast, H3K64Ac is enriched at actively transcribed regions (73), therefore its gradual increase during DDR observed at γH2AX mononucleosomes might promote local chromatin relaxation and DDR-induced histone exchange (Fig.6B, Extended Data Fig.6D). Taken together, we employed an innovative strategy to characterize dynamic profiles of 33 hPTMs. We distinguished DSB-induced kinetic profiles of hPTMs that occur globally from those marked by γH2AX, to causally associate hPTMs with DSB events at mono-nucleosome resolution.

### Integrative chromatin proteomics implicates G9A-mediated H3K56me1 in the DDR

The added value of our multi-layered approach resides in the integration of the complementary chromatin data to derive functional causality, and to infer novel aspects of the DSB repair. In particular, we profiled chromatin-associated abundance for different histone writers, erasers, and readers during DSB repair determined by iPOC (Fig.5G), and integrated this with kinetic profiles of their cognate DDR-induced and *foci*-specific hPTMs (Fig.6B). This shows that the chromatin recruitment for both writer and eraser of H3 and H4 acetylation (i.e. KAT7 and HDAC1, respectively) observed in iPOC, seems to be responsible for DDR-induced hPTM dynamics. In particular, decreased acetylation on H3 coincides with a reduced HDAC1 chromatin association, paralleled by the transient increase in both KAT7 and H4 di-acetylation (Fig.6C, Extended Data Fig.6D, E). Similarly, also H3K9me1 was globally induced at early time points of DNA repair, potentially as a consequence of the observed chromatin enrichment of EHMT2/G9A (Fig.6D, Extended Data Fig.6D). We could also link the kinetics of histone readers and their *foci*-specific hPTM, including the well-known crosstalk between dimethylated K20 on histone H4 (H4K20me2) and 53BP1, which drives towards NHEJ. In line with the replication-coupled dilution model (22), our findings suggest a rapid end joining-mediated repair of the DSBs, followed by a chromatin depletion of both these determinants at later time points of the repair process (Fig.6E, Extended Data Fig.6D). In addition, among the *foci*-specific hPTM, we observed a rapid increase of trimethylated K36 on histone H3 (H3K36me3) that is fully compatible with the chromatin enrichment observed in iPOC for SETMAR and PSIP1, which are the respective writer and reader of this modification. In contrast, the DSB-mediated chromatin recruitment of KDM2A seem to be decoupled from H3K36me3 (Fig.6F).

The identification of known crosstalk between chromatin regulators and their cognate hPTMs, prompted us to expand our investigation to unexpected pairing between such determinants. In particular, the DSB-mediated chromatin recruitment of G9A and the corresponding *foci*-specific monomethylation of its target residue K56 of histone H3 (H3K56me1) caught our attention (Fig.6G). H3K56 is located on the lateral surface of histone H3 close to DNA entry/exit site, and its acetylation seems to contribute to chromatin reassembly after DNA repair, while a role in cell cycle progression has been proposed for H3K56me1 (74, 75). Our data reveal that, in mononucleosomes flanking the break site, G9A-mediated monomethylation precedes the acetylation of H3K56 (1h, 4h) (Fig.6B, Extended Data Fig.6D, G), thus providing for the first time a role for H3K56me1 in the DDR. In the functional investigation of this novel epigenetic crosstalk, we examined and observed that inhibition of G9A with BIX-01294 dramatically decreased the efficiency of HR without perturbing cell cycle regulation (Fig.6H, Extended Data Fig.5N-Q). In line with these results, inhibition of G9A sensitized to treatment with PARPi alone or in combination with etoposide (Fig.6J). In particular, the selective inhibition of G9A did not affect RAD51 chromatin recruitment but resulted in a significant increase in BRCA1 loading at DSBs and deficient repair at these *foci* (Fig.6J, K). Moreover, the interaction between H3K56me1 and γH2AX is mediated by the induction of DSBs, and decreases upon treatment with low concentrations of G9Ai (Extended Data Fig.6F–H). Taken together, our results show that G9A is rapidly recruited to chromatin upon DSB, where it promotes monomethylation of H3K56 at γH2AX mononucleosomes. Since inhibition of G9A causes accumulation of BRCA1 *foci* and impairs homologous recombination repair, our data suggests that H3K56me1 may have a role in HR pathway, presumably by acting as docking site for proteins involved in HR repair downstream of BRCA1.

In summary, we showed how our multi-layered approach identified temporal dynamics shared between chromatin regulators and either cognate DDR-induced or DNA repair *foci*-specific modifications. Among the novel crosstalk identified, we further demonstrated the relationship between H3K56me1 and G9A in DNA damage, and showed that inhibition of this enzyme decreases HR repair efficiency and sensitizes BRCA-proficient cells to PARPi treatment, thus identifying a synthetic lethality with potential clinical relevance.

## DISCUSSION

Double-strand break repair takes place in the context of chromatin, where a coordinated mechanism engages histone PTMs and DNA-recruited regulators to set the stage for the efficient damage response cascade (6). To clarify this dynamic and composite picture, here we characterized the response to DSB from a chromatin-centric perspective using three exploratory proteomic-based approaches to investigate the chromatin organization at different levels of resolution, namely global chromatin composition (studied by iPOC), functional interaction maps of DDR core components (in ChIP-SICAP) (24), and hPTMs at monucleosomes flanking the break site (in N-ChroP) (26). Moreover, by adding a temporal dimension, we generated complementary data sets that collectively produce a comprehensive and detailed panorama of the chromatin dynamics during the DDR. This rich resource, which can be mined in an interactive fashion, led us to functionally characterize multiple proteins and hPTMs that had not been associated with DSB repair before, and assigned them a role in HR, NHEJ or pathway choice. In addition, for several unanticipated HR-associated proteins we identified synthetic lethality with PARPi in HR-proficient cells (HRDness or BRCAness), thus opening to clinically-relevant opportunities.

iPOC is a novel approach that we developed to capture, identify and quantify proteins recruited to or evicted from chromatin during the DDR. Salient features of iPOC include labeling of DNA by the thymidine analog EdU, as in iPOND and iPOTD (54, 76), combined with SILAC labeling for robust protein quantification, and capture of biotinylated complexes on protease-resistant streptavidin beads (25) to boost assay sensitivity. Differently from iPOND and NCC (55) that offer a site-specific view, the power of iPOC resides in its ability to determine compositional changes on a chromatin-wide scale, thereby complementing more targeted approaches like ChIP-SICAP. Indeed, beyond identifying core repair proteins also found in ChIP-SICAP, multiple proteins were exclusively identified in iPOC, notably proteins known for transient interactions like chromatin-modifying enzymes and transcriptional regulators (Fig.5G). This not only highlights the sensitivity of the approach, but also underscores the simultaneous recruitment of a functionally diverse set of proteins, and suggests that their respective biological activities cannot be seen in isolation. As illustrative example, we showed cross-regulation among functional processes through the negative modulation of HR mediated by the transcriptional regulator ADNP (Fig.5I). Here we applied iPOC to the DDR, however we envision that, in replicating cells, this approach can be extended to characterize changes in overall chromatin composition upon any molecular or cellular perturbation.

Another unique aspect of our study is the discrimination between DDR-induced and DNA repair *foci*-specific hPTMs by N-ChroP. Indeed, this approach brings together the unbiased nature of proteomics with the specificity of ChIP, to characterize hPTMs landscape of a specific chromatin region. In comparison with widely used ChIP-seq approaches (15), N-ChroP identifies cross-talks among hPTMs without *a priori* knowledge beside the modification of interest. Applied in a time-course fashion, this allowed us to confirm known and reveal novel PTM profiles, and to provide a refined view to published data. For instance, H3K79 acts as one of the docking sites for 53BP1 recruitment (66), however its methylation level has been reported to be unchanged or to decrease upon DSB (77) (15). Our dynamic data show that H3K79me2 follows a bi-modal increase at 0.5h and 4h from DSB induction (Fig.6B, Extended Data Fig.6D), while this modification is partially depleted at *foci*-specific level at 8h after IR, thus indicating that the time of sampling is crucial to explain these apparently conflicting observations. Similarly, H3K56Ac has been reported to be either deregulated or unchanged in the DDR (78). Our results may clarify these conflicting reports as we observed a *foci*-specific increase at late time points during DSB repair, accompanied by a decrease of this modification on a global scale (Fig.6B, Extended Data Fig.6D).

Since our proteomic approaches characterize complementary chromatin layers, their integration offers a detailed view to better understand the DNA repair process. Cross-correlation of these data may not prove direct causality, however they provide compelling examples of regulatory events such as between hPTMs and their cognate writers and readers. For instance, the overlay of N-ChroP and iPOC data indicate the cross-talk among the methyltransferase SETMAR, the reader PSIP and the *foci*-specific increase in trimethylated H3K36. Similarly we also identified coinciding temporal profiles of the hyper-acetylation of 9-17 peptide of histone H3 and the enrichment at chromatin for the histone acetyltransferase responsible for H3K14 acetylation KAT7, and of 53BP1 and dimethylated H4K20, a modification well-known to be involved in DSB repair regulation (21, 22) (Fig.5G, Fig.6B-F). Similarly, we identified the chromatin recruitment of G9A, an enzyme that can methylate H3K9 and H3K56. In our previous work, we could readily detect H3K9me2/3 (27), while we were unable to identify these hPTMs during the DDR. We therefore conclude that di- and tri-methylation of H3K9 occur at undetectable levels or do not correlate particularly with DNA damage, in agreement with recent data showing a lack of association between H3K9me2/3 and DSB repair (15). Interestingly however, we instead observed a *foci*-specific increase of monomethylated H3K56, another substrate of G9A, thus suggesting a functional link between G9A recruitment and H3K56me1. Moreover, induction of HRDness upon G9A inhibition (Fig.6H-I) in the absence of H3K9me2/3 suggests that, besides its function in DNA replication (75), H3K56me1 might have an as-yet unrecognized role in HR downstream of BRCA1. This illustrates how our complementary results represent a valuable resource to describe the complexity of chromatin response to DSBs (Fig.7) and to propose a possible mechanism of action for newly identified DSB-dependent chromatin proteins.

Through ChIP-SICAP and iPOC we identified novel candidates involved in the DDR, which we characterized by means of the traffic light reporter assay allowing for the monitoring of both NHEJ and HR events in the same cell population (79). This DSB cell system was fundamental especially to distinguish DDR components regulating both pathways (e.g. NuMA, SMARCE1 and PRPF8) from chromatin proteins belonging to HR (e.g. ATAD2, TPX2, G9A). Importantly, while both classes of proteins exhibit an overall decrease in HR efficiency, only the latter group shows sensitization to PARPi treatment (Fig.2B, E, Fig.4E, F, Fig.6H, I).

These findings demonstrate that our multi-layered chromatin approach powerfully complements genetic screens to identify HRDness, while potentially explaining the benefit of PARPi-therapy observed in BRCA wild-type patients (80) (81). In addition, newly identified chromatin regulators involved in DSB repair may be nominated as promising drug targets, spurring ongoing efforts to develop epigenetic drugs for targeted therapies or in combination with PARPi (82). Indeed, when looking for synthetic lethality, we could shortlist various targets to be investigated in knockdown screens; moreover the functional interaction identified *via* our chromatin-centered proteomics might mimic genetic interaction and suggest cooperativity between or among identified targets. Such data could also be a valuable resource for multigene perturbation technologies (83) to identify paralogues or functionally related proteins that may be a suitable target for such multiple knockdown approaches.

Our chromatin-directed analysis suffers from limitations in common with other proteomic-based strategies. First, these data do not indicate the chromatin conformation and the genomic localization of the described interactions. Second, iPOC and N-ChroP describe associations among chromatin determinants without discriminating between NHEJ-or HR-prone sites. Therefore, affinity-based enrichment of candidates coupled with sequencing represents the optimal orthogonal validation for both these aspects. Finally, so far we only explored a few classes of PTM and focused almost exclusively on histones without investigating modifications on non-histone proteins. For this reason, on the same line as previous studies (84), further multi-layered studies examining PTMs on other chromatin proteins will be complemental to our data.

In line with the idea of multi-omics integration (85), our findings show how multiple chromatin-centered approaches are needed to describe the complexity of a biological processes such as DNA damage repair. At the same time the temporal dimension is fundamental for defining the protein dynamics *via* unbiased approaches like mass spectrometry. To best of our knowledge, this strategy represents the first resource of chromatin-mediated functional interactions during DNA repair, and provides a focused yet comprehensive view of this biological process.

## Acknowledgments

The authors thank all members of the division Proteomics of Stem Cells and Cancer, German Cancer Research Center, Heidelberg for their support and advice. The authors thank Dr. Ali Bakr from the Division of Cancer Epigenomics, German Cancer Research Center (DKFZ) for discussion and for providing U2OS ID3-GFP cells. The authors also thank the DKFZ Light Microscopy Facility for its support. G. Legube (University of Toulouse, France) kindly supplied AID-DIvA cells, N. Ayoub (Technion, Israel) kindly provided U2OS-TLR cells, and R Syljuåsen (Oslo University Hospital, Norway) kindly provided the U2OS m53BP1-mCherry. TGH is supported by the Deutsche Forschungsgemeinschaft (DFG): SFB 1361 (Project-ID 393547839), project 19.

## Author contributions

JK and GS designed the research. TGH provided scientific expertise to the setup and early phase of the project. LA performed FACS-based experiment with resources and support from MS. GS and LA analyzed the data. NP realized the webpage, GS and JK wrote the manuscript with input from all authors.

## Declaration of Interest

The authors have no conflict of interest to declare.

## Additional resources

A webpage-based viewer is available at “chromatin-proteomics.dkfz.de”, where identified proteins and histone modifications can be searched and displayed in the chromatin layers where they are quantified.

Moreover, a dedicated tab shows dynamic trends of associated chromatin determinants (i.e. proteins and histone modifications), thus suggesting potential functional associations.

The ChIP-SICAP protocol is maintained at protocols.io: dx.doi.org/ 10.17504/protocols.io.bcrriv56.

## SUPPLEMENTAL FIGURE LEGENDS

**Extended Data Figure 1. Chromatin-mediated interactors quantified in RPA1, RAD50 and MDC1 ChIP-SICAP experiments. Related to Figure 1**. Log10 intensity-based quantification (iBAQ) rank distribution of ON-chromatin functional interactors quantified in RPA (**A**), RAD50 (**B**), MDC1 (**C**) ChIP-SICAP experiment. Light (L), medium (M) and heavy (H) SILAC channels correspond to technical control, untreated condition (UT), and 1h upon IR-induced DSBs (IR), respectively. Histone proteins are highlighted in blue, protein used as bait in ChIP is reported in red, examples of core components of HR and NHEJ are shown in black. Venn diagrams show the overlap between proteins quantified in UT and IR condition, while stacked histograms represent the number of interactors identified exclusively in the ON-chromatin fraction (ON-chromatin only) or constitutively associated with the target (Chromatin) in RPA (**D**), RAD50 (**E**), or MDC1 (**F**) experiment. Tables show ON-chromatin functional interactors quantified exclusively in IR condition. ChIP-SICAP experiment performed in UT (top) or IR condition (middle) and DSB-induced modulation of ON-chromatin interactors (bottom) of 53BP1 (**G**) or XRCC6 (**H**). Scatterplots show the correlation between IR-induced interactors of 53BP1 (**I**), XRCC6 (**J**), or MDC1 (**K**), while the bait proteins are in their OFF-(x-axis) or ON-chromatin (y-axis) fraction. Average values of biological replicates are shown. Ctr = isogenic control experiment. Values into parenthesis correspond to log2 IR/Ctr ratios in the two chromatin fractions. n.d. = protein not identified in that experiment. imp = protein not detected in Ctr, value replaced with mean of log2 ratio distribution. Histone proteins are shown into circle.

**Extended Data Figure 2. Functional characterization of targets interacting on-chromatin with HR and NHEJ core components. Related to Figure 2. A**) Western blot validation of siRNA-mediated silencing of targets shown in figure 2. Mk = protein ladder; Inp = input, red arrows show protein target band. **B**) Representative flow plots of U2OS-TLR cell lines following transduction with I-SceI-T2A-IFP, GFP-donor-BFP plasmids and silenced for ON-chromatin targets related to figure 2. TLR readout is shown following gating on live single cells double positive for IFP and BFP. Cells knockdown for 53BP1 and RPA/RAD51 are shown as positive controls for NHEJ and HR impairment, respectively. **C**) Quantification of EGFP (HR) and mCherry (NHEJ) cells as percentage of double positive cells in 3 independent experiments, with standard deviation. **D**) Plot of PI-based flow cytometry analysis of cell cycle in U2OS cells knockdown for targets related to figure 2. After gating on live cells, the percentage of cells in G1, S and G2-M phases was determined and reported in stacked histogram with standard deviation among biological replicates (**E**). siCtr = isogenic siRNA control experiment. Candidate-centered boxplot representation of RAD51 (**F**) and BRCA1 (**G**) *foci* during DSB repair in AID-DIvA cells silenced for the reported proteins related to figure 2. * and ** correspond to p-value < 0.05 and 0.01 of ANOVA statistical test, respectively.

**Extended Data Figure 3. Dynamics of γH2AX-specific interactors during DNA response. Related to Figure 3**. Density plot of log2 SILAC ratio distribution between γH2AX and technical IgG isogenic control (left, H/L) in untreated condition (**A**), or 0.5h (**B**), 1h (**C**), 4h (**D**), 8h (**E**) upon DSB induction. Two replicates are shown per condition. Scatterplots show the correlation between log2 H/M SILAC ratios (γH2AX/H2A) in replicates at specified time point. The positive control MDC1 is highlighted in red, while protein targets related to figure 3 (THRAP3, ATAD2 and TPX2) are reported in black in the scatterplot. Proteins identified as important for HR and NHEJ in figure 2 are shown in green and orange, respectively. **F**) Bar plot represents the frequency of γH2AX-specific interactors involved in the listed categories. **G**) Functional classification of γH2AX-specific interactors already reported to be involved the DDR. Functional protein network visualization and GO categories associated with proteins rapidly recruited (**H**) or evicted (**I**) from γH2AX shown in figure 3.

**Extended Data Figure 4. Functional characterization of targets interacting with γH2AX during DNA damage repair. Related to Figure 4. A**) Validation of THRAP3 antibody specificity *via* immunofluorescence in proficient or siTHRAP3 cells. **B**) Western blot validation of siRNA-mediated silencing of TPX2 and ATD2 shown in figure 4. **C**) Candidate-centered boxplot representation of γH2AX *foci* during DSB repair in AID-DIvA cells silenced for the reported proteins related to figure 4. **D**) Representative flow plots of U2OS-TLR cell lines silenced for ON-chromatin targets related to figure 4. Cells knockdown for 53BP1 and RPA/RAD51 are shown as positive controls for NHEJ and HR impairment, respectively. **E**) Quantification of EGFP (HR) and mCherry (NHEJ) cells as percentage of double positive cells in 3 independent experiments, with standard deviation. **F**) Representative PI-based flow cytometry analysis of cell cycle in U2OS cells knockdown for TPX2 or ATAD2. The percentage of cells in G1, S and G2-M phases is reported in stacked histogram with standard deviation among three biological replicates (**G**). Candidate-centered boxplot representation of RAD51 (**H**) and BRCA1 (**I**) *foci* during DSB repair in AID-DIvA cells silenced for the reported proteins related to figure 4. *, ** and *** correspond to p-value < 0.05, 0.01 and 0.001 of ANOVA statistical test, respectively.

**Extended Data Figure 5. Setup of iPOC in DNA damage repair, comparison with fractionated chromatin input and characterization of the functional role of regulators rapidly recruited at chromatin during DNA damage repair. Related to Figure 5 and 6. A**) EdU-based DNA labeling specificity in cells untreated (top) or upon 1h from IR (bottom) exposed to increasing amounts of EdU for 18h and subjected to CuAAC click chemistry with Cy5-azide fluorophore in comparison with cells not treated with EdU (left, NO EdU). Pictures on the far right refer to cells exposed to 20µM EdU for 18h, subjected to CuAAC click chemistry-based DNA labeling with biotin-azide and visualized with Alexa488-(green) or Alexa594-(red) conjugated anti-biotin antibodies. **B**) Number of γH2AX *foci* per nucleus in U2OS cells exposed for 18h to increasing concentrations of EdU (0µM to 200µM) and either left untreated or subjected to ionizing radiations (5Gy, 1h recovery). p-value from two-sided t-test statistics are reported. n.s. = not statistically significant. **C**) Multi scatterplots showing the correlation of log2 SILAC ratios for proteins quantified in iPOC at untreated condition (medium) over the light negative control (UT/Ctr). Venn diagram shows the overlap of quantified proteins. Pearson correlation coefficient is shown in blue. **D**) As in C) but for determinants enriched in heavy-labeled samples at 1h, 4h and 8h time points upon DSB induction over the light negative control (Ctr). **E**) Correlation between heavy-labeled and medium-labeled untreated sample at different time points during the DDR. Pearson and Spearman correlation coefficients are shown in blue and red, respectively. **F**) Triple SILAC-based experimental design for chromatin input deep investigation during the DDR. Triple SILAC labeled U2OS cells were collected at 0.5h, 1h, 4h, or 8h upon 5Gy ionizing radiation-induced DSBs and combined in equal amounts into triple SILAC experiments. Extracted chromatin was sheared, subjected to tryptic digestion and high pH peptide fractionation prior MS analysis. **G**) Multi scatterplot shows log2 SILAC ratios for proteins quantified at different time points over untreated (UT) condition. For each pair, Pearson correlation coefficient and number of quantified proteins are reported in blue and red, respectively. **H**) Top5 gene ontology categories associated with proteins deregulated in the chromatin input at different time points during the DDR. **I**) Heatmap representation of protein abundance changes during DSB repair quantified through fractionated chromatin input analysis. Average log2 SILAC ratios are shown upon z-score normalization. **J**) Heatmap representation of SILAC log2 ratios for proteins quantified in iPOC at different time points during DSB repair over technical control (Ctr) or untreated sample (UT) in comparison with the protein dynamics quantified in the chromatin input (Input over UT). Red arrows highlight PHF14, G9A, SMARCA1 and ADNP proteins shown in figure 5. **K**) Average log2 SILAC ratio over untreated control for different histone variants quantified in iPOC (blue scale) and in chromatin input (ChrInp, orange scale). Histone variants not identified are indicated with nd. **L**) Western blot validation of siRNA-mediated silencing of targets shown in shown in figure 5. **M**) Candidate-centered boxplot representation of γH2AX *foci* during DSB repair in AID-DIvA cells silenced for the reported proteins related to figure 5. **N**) Representative flow plots of U2OS-TLR cell lines silenced for PHF14, ADNP and SMARCA1 or upon G9A drug inhibition (G9Ai). Cells knockdown for 53BP1 and RPA/RAD51 are shown as positive controls for NHEJ and HR impairment, respectively. **O**) Quantification of EGFP (HR) and mCherry (NHEJ) cells as percentage of double positive cells in 3 independent experiments, with standard deviation. **P**) Plot of PI-based flow cytometry analysis of cell cycle in U2OS cells knockdown for PHF14, ADNP and SMARCA1 or upon G9A inhibition. The percentage of cells in G1, S and G2-M phases is reported in stacked histogram with standard deviation among biological replicates (**Q**). Candidate-centered boxplot representation of RAD51 (**R**) and BRCA1 (**S**) *foci* during DSB repair in AID-DIvA cells silenced for the reported proteins related to figure 5 and 6. ** and *** correspond to p-value < 0.01 and 0.001 of ANOVA statistical test, respectively.

**Figure 5.**
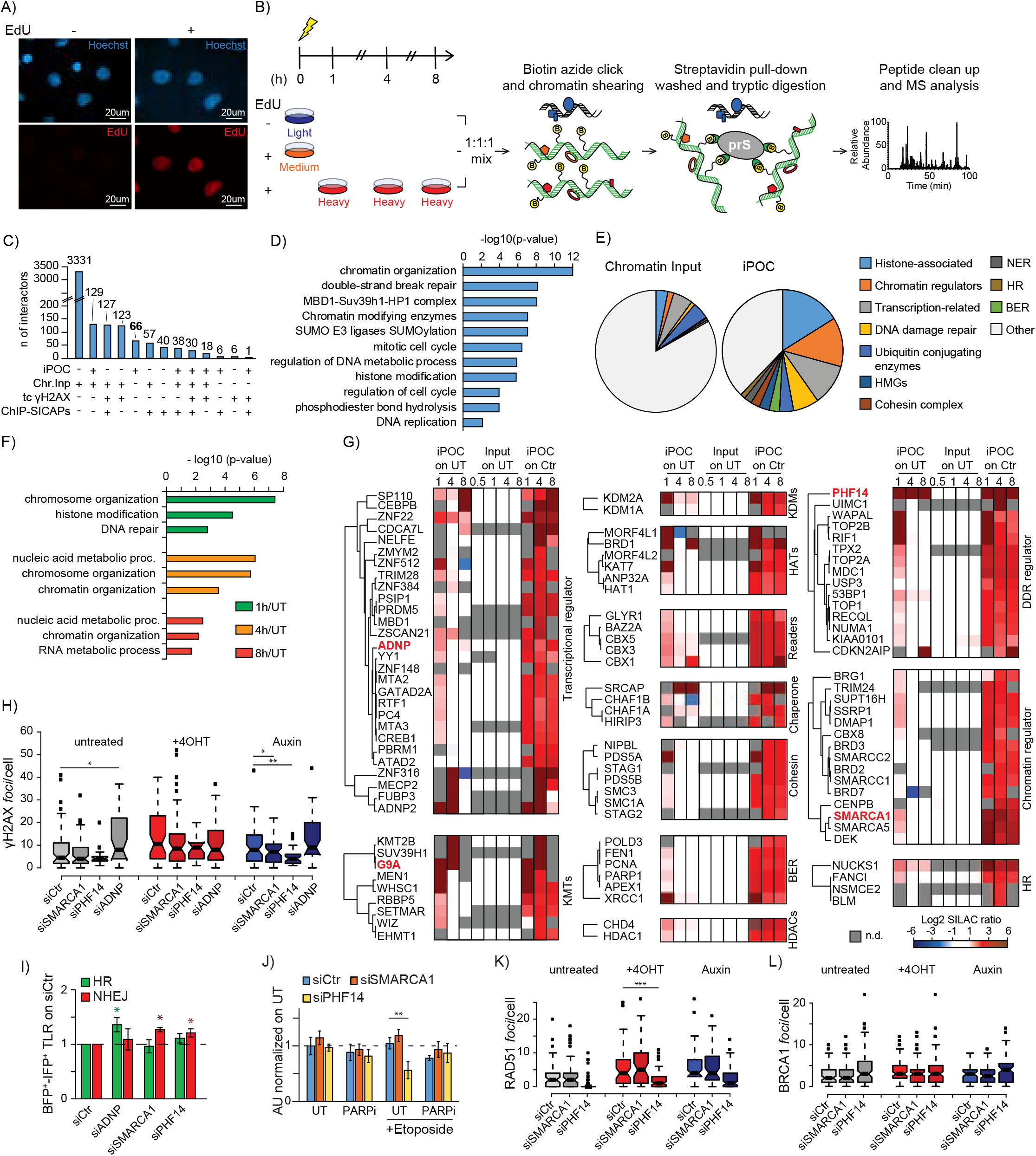
Chromatin composition dynamics during DDR investigated by iPOC. **A**) Representative immunofluorescence of cells treated with DMSO (left) or the nucleotide analogue EdU (right), and subjected to click chemistry with Alexa594-azide fluorophore. **B**) Schematic representation of iPOC experimental strategy. SILAC labeled U2OS cells are exposed to either DMSO (light, as control) or EdU (medium and heavy) and crosslinked in untreated condition or at different time points during the DDR. Cells are then subjected to permeabilization, followed by click chemistry-based DNA labeling with biotin azide and enrichment with protease-resistant streptavidin beads (prS) prior to stringent washes, tryptic digestion and MS-based identification of chromatin-associated proteins. **C**) Bar chart displays the overlap of functional interactors identified in ChIP-SICAP as from figure 1 (ChIP-SICAPs), in time-course γH2AX-experiment from figure 3 (tc γH2AX), in iPOC and in fractionated chromatin preparation used as input for ChIP (Chr.Inp). **D**) Top 11 gene ontology categories associated with targets exclusively quantified in iPOC. **E**) Comparison between the frequency of listed categories of protein in chromatin input (left) and in iPOC (right). **F**) Gene ontology (GO) categories associated with proteins identified in iPOC as recruited at chromatin at the different time points upon ionizing radiation (IR) compared with untreated condition. **G**) Hierarchical clustering heatmap representation of SILAC ratios (in log2) for functional categories of DDR- and chromatin-regulators quantified in iPOC at different time points during DSB repair over technical control (Ctr) or untreated sample (UT). For all targets, the relative dynamics quantified in the chromatin input over the respective untreated condition (Input on UT) is also shown. Proteins further followed up are highlighted in red. n.d.: not detected. **H**) Number of γH2AX *foci* per nucleus in AID-DIvA cells after target knockdown upon DSB induction and repair (tamoxifen/+4OHT and Auxin, respectively), in comparison with on-target non-targeting silencing control (siCtr). **I**) Quantification of HR (green) and NHEJ (red) repair events in U2OS-TLR cells depleted for ADNP, SMARCA1, or PHF14 and normalized on silencing control (siCtr). Green and red asterisks reflect significant regulation in HR and NHEJ, respectively, from siCtr. **J**) Quantification of CFA in U2OS cells depleted of PHF14 or SMARCA1 and subjected to PARP inhibitor (PARPi) alone or in combination with etoposide to promote DSBs formation. Mean values of three replicates normalized to untreated condition (UT), error bars represent standard deviations. Quantification of RAD51 (**K**) and BRCA1 (**L**) *foci* in AID-DIvA cells after target knockdown. *, ** and *** correspond to p-value < 0.05, 0.01 and 0.001 of ANOVA statistical test, respectively.

**Figure 6.**
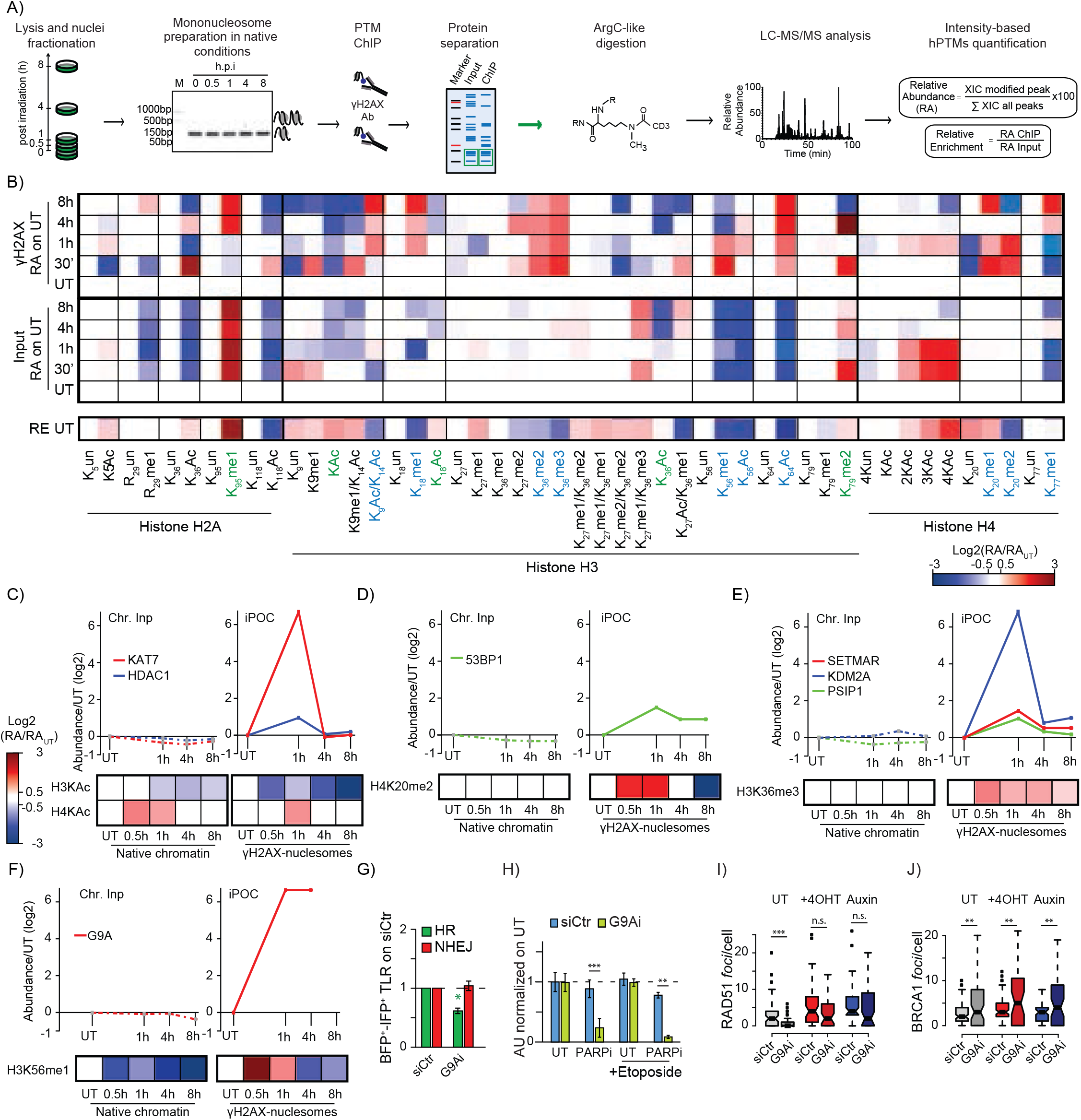
Profiling of hPTMs during DDR at mononucleosome resolution. **A**) Experimental design of time course N-ChroP. Mononucleosomes from cells at different time points during the DDR (hour post irradiation) are enriched for γH2AX and associated hPTMs are quantified *via* MS upon chemical derivatization and tryptic digestion (ArgC-like digestion). For each hPTM the relative abundance (RA) is calculated as the extracted ion chromatography (XIC) of a specific modification over the sum of XICs for all modifications occurring on the same peptide. The relative enrichment (RE) of each hPTM represents the ratio between its RA in the enriched ChIP over the RA in the mononucleosome input. **B**) Heatmap representation of the log2 RA for hPTMs in γH2AX-enriched mononucleosomes (γH2AX) or in native mononucleosome preparation used as input (Input) calculated over the respective untreated sample (UT). Relative enrichment at steady state (RE UT) highlights the specific combinatorial patterns of hPTMs at γH2AX-marked mononucleosomes. DDR-induced and *foci*-specific modifications are highlighted in green and light blue, respectively. Abundance relative to UT conditions for writers (red), erasers (blue), or readers (green) associated with H3/H4 acetylation (**C**), H4K20me2 (**D**), H3K36me3 (**E**), and H3K56me1 (**F**). Lines correspond to protein abundance in chromatin input (Chr.Inp, left) or in iPOC (right). Heatmap represents RA of hPTM during DSB repair in the native chromatin input (Native chromatin, left) or in γH2AX-marked mononucleosomes (γH2AX-mononucleosomes, right). **G**) Quantification of HR (green) and NHEJ (red) repair events in U2OS-TLR cells upon inhibition of G9A (G9Ai) normalized to silencing control (siCtr). Green asterisk reflects significant regulation in HR compared with siCtr. **H**) Quantification of CFA in U2OS cells upon G9A inhibition and subjected to PARP inhibitor (PARPi) alone or in combination with etoposide treatment. Mean values normalized to untreated condition (UT) and standard deviations are shown. Quantification of RAD51 (**I**) and BRCA1 (**J**) *foci* in AID-DIvA cells upon drug-inhibition of G9A (G9Ai). ** and *** correspond to p-value < 0.01 and 0.001 of ANOVA statistical test, respectively. n.s. = not significant. UT = untreated conditions, +4OHT = upon DSB formation, Auxin = resolution of DNA breaks.

**Figure 7.**
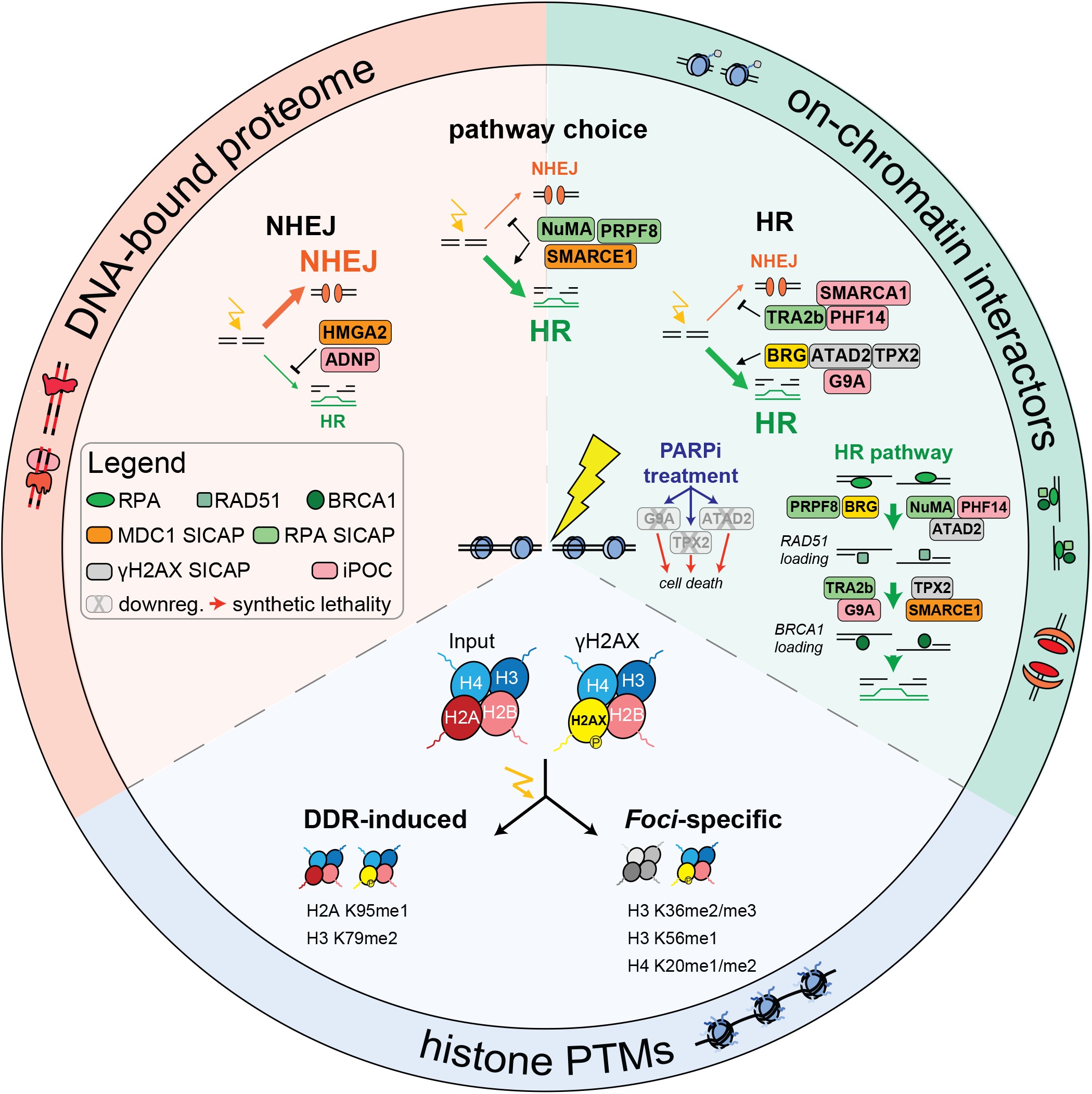
Multi-layered chromatin-directed proteomics identifies cell vulnerabilities in HR and NHEJ. Integration of orthogonal approaches in chromatin-directed proteomics identifies chromatin proteins contributing to NHEJ, repair pathway choice, and HR; moreover it distinguishes trends of hPTMs induced globally during the DNA damage repair (DDR induced) from modifications specifically enriched at γH2AX-mononucleosomes (*foci*-specific). Orthogonal imaging-based strategies determine the role in the early steps of homologous recombination repair (HR pathway) for chromatin targets involved in either HR or pathway choice, while CFA assays identify targets whose downregulation drives towards synthetic lethality with PARPi epigenetic drug (PARPi treatment).

**Extended Data Figure 6. Setup of native enrichment of γH2AX-marked mononucleosomes and dynamics of co-associated histone modifications during DNA damage response. Related to Figure 6. A**) Time course MNase digested chromatin to mononucleosome preparation from cells in untreated condition or at the indicated time points during the DDR. **B**) Coomassie stained SDS-PAGE of chromatin input and γH2AX-enriched mononucleosomes (ChIP) at the different time points during DSB repair. Log2 RA dynamics of hPTMs on histone H2A (**C**), H3 (**D**) or H4 (**E**) in either native chromatin input (Input, up) or in γH2AX-enriched mononucleosomes (γH2AX ChIP, down). Mean values are shown. Respective stack histograms represent the percentage of RA (RA%) for each hPTM in either native chromatin input (Inp) or in γH2AX mononucleosomes (γH2AX) at different time points during the DDR. **F**) Mononucleosome preparation from U2OS cells untreated or treated with G9Ai or Etoposide (Etop), alone or in combination. **G**) H3K56me1 ChIP from mononucleosomes preparation shown in (F) and western blot analysis of H3K56me1 and γH2AX (red arrow), total H3 is used as loading control. G9Ai inhibition corresponds to 5µM for 4h. **H**) Western blot analysis of H3K56me1 (top) and H3K27me3 (bottom) levels in cells left untreated or treated for 4h with increasing concentrations of either G9Ai or EZH2i. Quantification refers to H3K56me1 over H3K27me3 signal normalized on UT conditions.

## Resource availability

### Lead contact

Further information and requests for resources and reagents should be directed to and will be fulfilled by the lead contact, Prof. Jeroen Krijgsveld

### Materials availability

This study did not generate new unique reagents

### Data and code availability

The webpage-based viewer for the experiments described in this manuscript is available at “chromatin-proteomics.dkfz.de”.

The mass spectrometry proteomics data have been deposited to the ProteomeXchange Consortium *via* the PRIDE (https://www.ebi.ac.uk/pride/) (86) partner repository with the dataset identifier PXD027421.

### Experimental model and subject details

U2OS (ATCC) and U2OS-ID3-GFP (kindly provided by Dr. Ali Bakr, DKFZ, Heidelberg) cell lines were cultured in Dulbecco’s modified Eagle’s medium (DMEM) supplemented with antibiotics, and 10% FCS (Invitrogen) at 37°C under a humidified atmosphere with 5% CO_2_. U2OS-TLR cells (kindly provided by Prof. Ayoub N) were cultured as U2OS with puromycin 0.6µg/mL. U2OS pIRES-mCherry-m53BP1 (kindly provided by Prof. Randi Syljuåsen, Oslo University Hospital) and AID-DIvA (MTA with Dr. Gaelle Legube, CBI, Toulouse) were cultured as U2OS with 800µg/mL G418, Geneticin (Thermo Fisher Scientific, 10131035). DSB were induced in AID-DIvA with 300nM hydroxyl tamoxifen (Sigma Aldrich, T5648) for 4h. Repair was promoted with 500µg/mL Auxin (Indole-3-acetic acid sodium salt, Sigma Aldrich, I5148) for 1h.

## Method details

### Chromatin-associated interactors investigated through ChIP-SICAP

ChIP-SICAP experiments were performed as described before (24) with some modifications. In brief, U2OS cells were metabolically labeled in SILAC (Ong SE et al 2002) DMEM medium containing light (Arg0, Lys0), medium (Arg6, Lys4) or heavy (Arg10, Lys8) amino acids. For experimental design related to figure 1, U2OS cell pellets corresponding to 24 × 10^6^ cells were cross-linked with formaldehyde (1% final concentration) in the absence (medium) or after 1h recovery from 5Gy ionizing radiation (heavy) elicited with Gammacell 40 Exactor (Best Theratronics). Control sample labeled in light corresponds to a 1:1 mix of 12 × 10^6^ untreated cells and 12 × 10^6^ irradiated cells as above. Cell pellets were resuspended in 5mL of lysis buffer 1 (50mM HEPES-KOH pH 7.5, 140mM NaCl, 1mM EDTA, 10% Glycerol, 0.5% NP-40, 0.25% Triton X-100), rotated on the wheel at 4°C for 10min and spun at 400g for 5min at 4°C. Pellets were then resuspended in 5mL of lysis buffer 2 (10mM Tris-HCl pH 7.5, 200mM NaCl, 1mM EDTA), rotated for 10min on the wheel at RT and spun at 400g for 5min at RT. Pellets were resuspended in 1.8mL of lysis buffer 3 (10mM Tris-HCl pH 7.5, 100mM NaCl, 1mM EDTA) and split in 6x 1.5mL sonication tubes (Diagenode). Upon 8 cycles of sonication with Pico Bioruptor (Diagenode) (30sec ON/ 30sec OFF), Triton X-100 (1% final concentration) was added to the samples and the tubes were spun at 400g. Supernatants from the same SILAC labeling were pooled. The supernatant from medium and heavy samples were combined in equal amounts and subjected to ChIP against target protein (between 5 and 15µg of antibody depending on the target), light-labeled controls were subjected to ChIP with isogenic IgG and used as internal control. After overnight incubation in the cold room, ProteinG magnetic beads previously coated overnight with BSA 0.5% in PBS1x (100uL per 10µg of antibody) were added. After 3h of rotating in the cold room, the beads were cleaned up with Tris–HCl 10mM. Next, the beads were treated with terminal deoxynucleotidyl transferase (EP0162) and biotinylated nucleotides (Biotin-11-dCTP, Jena Bioscience). The beads were then washed with IP buffer (50mM Tris–HCl pH 7.5, 1% Triton X-100, 0.5% NP-40, 5mM EDTA), and proteins were eluted with elution buffer (7.5% SDS, 200mM DTT) for 15 min at 37°C. Eluted samples were diluted in IP buffer and combined. Then, 100µL of protease-resistant streptavidin (or prS (25)) beads were added for the DNA enrichment. To analyze the soluble interactors, supernatant was concentrated with speedvac, subjected to SP3 protein clean up as previously described (87, 88) and eluted in AmBic 50mM prior digestion with 300ng trypsin (Promega V5280). For the on-chromatin interactors, prS beads were washed three times with SDS washing buffer (10mM Tris–HCl pH 8, 1% SDS, 200mM NaCl, 1mM EDTA), once with BW2x buffer (10mM Tris–HCl pH 8, 0.1% Triton X-100, 2M NaCl, 1mM EDTA), once with isopropanol 20% in water, and three times with acetonitrile 40% in water. The beads were transferred to PCR tubes using acetonitrile 40%. The supernatant was removed, and the beads were resuspended in 15µL of 50mM AmBic-10mM DTT. Then, the samples were incubated at 50°C for 15min to reduce the disulfide bonds. The cysteines were alkylated with 20mM IAA final concentration for 15min in the dark. IAA was neutralized by adding 10mM DTT final concentration. To digest the proteins, 300ng of trypsin (Promega V5280) was added to each tube. The digestion was performed for 18h and peptides were cleaned using SP3 beads and eluted in 0.1% trifluoroacetic acid (TFA) before mass spectrometry analysis.

For the experimental design shown in figure 3, light-, medium-and heavy-labeled cells were collected at untreated conditions or after 30min, 1h, 4h, 8h upon 5Gy IR. At each time-point, light-labeled sample was used for ChIP with isogenic IgG, while medium- and heavy-labeled samples were subjected to H2A and γH2AX ChIP (10µg of antibody). Samples were then mixed in equal amounts before DNA labeling. The rest of the procedure was carried out as described above.

### Isolation of Protein on Chromatin (or iPOC) during double-strand break repair

Per each experiment, 40× 10^6^ U2OS cells were metabolically labeled in SILAC (Ong SE et al 2002) DMEM medium containing light (Arg0, Lys0), medium (Arg6, Lys4) or heavy (Arg10, Lys8) amino acids. Medium- and heavy-labeled samples were also treated with 5-ethynyl-2’-deoxyuridine (or EdU) for 18h at a final concentration of 20µM. Light- and medium-labeled samples were collected in untreated conditions, while heavy labeled cells were harvested at 1h, 4h or 8h upon 5Gy ionizing radiations. Each cell pellet was resuspended in 4mL of permeabilization buffer (0.25% Triton X-100 in PBS1x), incubated for 30min at RT and spun at 4°C for 5min at 900g. Pellets were washed once with cold 0.5% BSA in PBS 1x and once with PBS 1x. Upon centrifugation as above, each pellet was resuspended in 5mL of click reaction mix (10µM biotin azide, Jena Bioscience CLK-047), 10mM sodium ascorbate, 2mM CuSO_4_ in PBS 1x) and rotated on the wheel, at RT, in the dark for 3h. Samples were then spun and washed once with 0.5% BSA in PBS1x and once with PBS 1x as above. The pellet was resuspended in 400µL of lysis buffer (1% SDS, 50mM Tris-HCl pH 7.5) and sonicated with the Pico Bioruptor (Diagenode) for 20 cycles (30sec ON/ 30sec OFF) or until a chromatin input focused around 600-700bp. Differently labeled samples were then spun at RT for 10min at 16000g, and supernatants were combined in equal amount before adding 200µL of magnetic prS beads (25) pre-conditioned with lysis buffer. Samples were rotated overnight on the wheel. Supernatant was discarded and prS beads-chromatin sample complexes were recovered on the magnet, washed once with lysis buffer and once with 1M NaCl. PrS beads-chromatin sample complexes where conditioned in AmBic 50mM, then resuspended in 30uL of 50mM AmBic-10mM DTT. Then, the samples were incubated at 50°C for 15min to reduce disulfide bonds, followed by cysteines alkylation with 20mM IAA for 15min in the dark. IAA was neutralized by adding 10mM DTT final concentration and proteins were digested with 300ng of trypsin (Promega V5280) for 18h. Peptides were cleaned using SP3 protocol as previously described (87, 88), and peptides were then eluted in 0.1% trifluoroacetic acid (TFA). Ammonium formate 20mM final concentration was added to each sample before subjecting them to fractionation using high-pH reverse-phase chromatography. Peptides were fractionated on an Agilent 1200 Infinity HPLC system with a Gemini C18 column (3µm, 110Å, 100 × 1.0mm, Phenomenex) using a linear 60min gradient from 0% to 35% (v/v) acetonitrile in 20mM ammonium formate (pH 10) at a flow rate of 0.1ml/min. Elution of peptides was detected with a variable-wavelength UV detector set to 254nm. Thirty-two 1-min fractions were collected and subsequently pooled into four fractions per sample.

### Chromatin preparation and fractionation to study the global dynamics of determinants during DNA repair

U2OS cells were metabolically labeled in SILAC (Ong SE et al 2002) DMEM medium containing light (Arg0, Lys0), medium (Arg6, Lys4) or heavy (Arg10, Lys8) amino acids. Each cell pellet corresponded to 24 × 10^6^ cells cross-linked with formaldehyde 1% final concentration. Medium-labeled samples were collected after either 30min or 4h from DSB induction with 5Gy ionizing radiations (IR) with Gammacell 40 Exactor (Best Theratronics), while heavy-labeled cells were harvested after either 1h or 8h from IR. Cells labeled with light amino acids were collected in untreated conditions and used as common reference channel between the two triple SILAC experiments. The first SILAC experiment contained cells untreated, 30min and 1h upon IR, labeled in light, medium and heavy channel, respectively. In the second experiment cells untreated, 4h and 8h upon IR were labeled in light, medium and heavy, respectively. Cell pellets from the same experiment were resuspended in lysis buffer 1 (50mM HEPES-KOH pH 7.5, 140mM NaCl, 1mM EDTA, 10% Glycerol, 0.5% NP-40, 0.25% Triton X-100) and mixed in equal amounts into the two triple SILAC experiments. Cells were then rotated on the wheel at 4°C for 10min and spun at 400g for 5min at 4°C, ultimately resuspended in lysis buffer 2 (10mM Tris-HCl pH 7.5, 200mM NaCl, 1mM EDTA). Upon rotation for 10min on the wheel at RT and centrifugation at 400g for 5min at RT, the pellets were resuspended in lysis buffer 3 (10mM Tris-HCl pH 7.5, 100mM NaCl, 1mM EDTA) and subjected to 8 cycles of sonication with Pico Bioruptor (Diagenode) (30sec ON/ 30sec OFF). Triton X-100 1% final concentration was added to the samples and the tubes were spun at 400g. Per each experiment, 20µg of chromatin was subjected to buffer exchange through SP3 protein clean up protocol (87) and resuspended in AmBic-10mM DTT before proteins digestion with 1:50 trypsin (Promega V5280) for 18h. Peptides were cleaned using SP3 beads and eluted in 0.1% trifluoroacetic acid (TFA) plus 20mM ammonium formate pH10 final concentration before high pH reverse-phase chromatography fractionation. Peptides were fractionated on an Agilent 1200 Infinity HPLC system with a Gemini C18 column (3µm, 110Å, 100 × 1.0mm, Phenomenex) using a linear 60min gradient from 0% to 35% (v/v) acetonitrile in 20mM ammonium formate (pH 10) at a flow rate of 0.1ml/min. Elution of peptides was detected with a variable wavelength UV detector set to 254nm. Forty 1-min fractions were collected and subsequently pooled into 8 fractions per sample experiment.

### Profiling of histone post-translational modifications during DNA repair in native conditions and at mononucleosome resolution

Native Chromatin Proteomic (or N-ChroP) protocol was performed as described before (26, 27) with some modifications. Fifty million U2OS cells were collected in untreated conditions or 30min, 1h, 4h, 8h recovery from 5Gy ionizing radiations elicited with Gammacell 40 Exactor (Best Theratronics) and resuspended in Native lysis buffer (10% sucrose, 0.5mM EGTA pH 8.0, 15mM NaCl, 60mM KCl, 15mM HEPES, 0.5% Triton, 0.5mM PMSF, 1mM DTT, 5mM NAF, 5mM Na_3_VO_4_, 5mM NaButyrate), supplemented with Triton X-100 0.5% final concentration and incubated for 10min on the wheel at 4°C. Nuclei were isolated *via* centrifugation on a sucrose cushion (Native lysis buffer with 20% sucrose) at 2800g for 30min, at 4°C. Pelleted nuclei were washed twice with PBS 1x at RT and resuspended in MNase digestion buffer (0.32 M sucrose, 50mM Tris-HCl pH 7.5, 4mM MgCl_2_, 1mM CaCl_2_, 0.1mM PMSF) digested to mononucleosomes with 0.01U/µL micrococcal nuclease (New England Biolabs, MO247S) at 37°C and spun at 2800g for 10min at 4°C. After saving 30µg as input, the supernatant was used for ChIP with γH2AX overnight, at 4°C on the wheel. ProteinG magnetic beads, pre-conditioned with BSA 0.5% in PBS1x (100uL per 10µg of antibody), were then added and left rotating for 3h at 4°C. Beads were then recovered on the magnet and washed twice with native washing buffer A (or WBA: 50mM Tris-HCl pH 7.5, 75mM NaCl, 10mM EDTA), once with native washing buffer B (WBA with 125mM NaCl) and once with native washing buffer C (WBA with 175mM NaCl). Immunopurified material and respective input were separated by SDS-PAGE. After Coomassie staining, histone bands were excised from the gel, and de-stained with 6 washes of alternate 0, 50%, 100% acetonitrile. Histones were then chemically alkylated with D6-acetic anhydride (Sigma 175641) in 1M AmBic for 3h at 37°C, followed by acetonitrile washes as above and in-gel tryptic digestion. Peptides were extracted from the gel with 3x 100% acetonitrile, 1x 5% formic acid, 2x 100% acetonitrile washes. Supernatants containing peptides were pooled, concentrated with speedvac and desalted with self-made StageTips (89) with C_18_ resin. Eluted peptides were lyophilized, resuspended in 0.1% TFA and analyzed with Q-Exactive HF (Thermo Scientific) mass spectrometer. Acquisition details under “Mass spectrometry data acquisition”.

### Mass spectrometry data acquisition

Peptides were loaded on a trap column (PepMap100 C18 Nano-Trap 100µm × 20mm) and separated over a 25cm analytical column (Waters nanoEase BEH, 75μm × 250mm, C18, 1.7μm, 130Å) using the Thermo Easy nLC 1200 nanospray source (Thermo Easy nLC 1200, Thermo Fisher Scientific). Solvent A was water with 0.1% formic acid and solvent B was 80% acetonitrile, 0.1% formic acid. During the elution step, the percentage of solvent B increased in a linear fashion from 3 to 8% in 4min, then increased to 10% in 2min, to 32% in 68min, to 50% in 12min and finally to 100% in a further 1min and went down to 3% for the last 11 min. Peptides were analyzed on a Tri-Hybrid Orbitrap Fusion mass spectrometer (Thermo Fisher Scientific) operated in positive (+2.5 kV) data dependent acquisition mode with HCD fragmentation. The MS1 and MS2 scans were acquired in the Orbitrap and ion trap, respectively, with a total cycle time of 3s. MS1 detection occurred at 120,000 resolution, AGC target 1E6, maximal injection time 50ms and a scan range of 375–1500 m/z. Peptides with charge states 2–4 were selected for fragmentation with an exclusion duration of 40s. MS2 occurred with NCE 33%, detection in topN mode and scan rate was set to Rapid. AGC target was 1E4 and maximal injection time allowed of 50ms. Data were recorded in centroid mode.

For histone samples, peptides were loaded and separated on same trap and analytical columns as above, but the percentage of solvent B increased in a linear fashion from 8 to 40% in 100min, then increased to 60% in 3min, to 95% in 5min and remained 95% for 3min before going back to 8% for 5min. Peptides were analyzed on a Q-Exactive HF Orbitrap mass spectrometer (ThermoFisher Scientific) operated in positive (+2.5kV) data dependent acquisition mode with HCD fragmentation. MS1 detection occurred at 60,000 resolution, AGC target 3E6 maximal injection time 150ms and a scan range of 300– 1500m/z. Peptides with charge states 2–5 were selected for fragmentation with an exclusion duration of 60s. MS2 occurred with NCE 30%, detection in top20 mode. AGC target was 5E4 and maximal injection time allowed of 250ms. Data were recorded in centroid mode.

### MS data processing, analysis and data visualization

Mass spectrometry data were processed with MaxQuant software (1.5.2.8, 1.6.2.6) (90, 91) using default settings. MSMS spectra were searched against the Human UniProt database concatenated to a database containing protein sequences of contaminants. Enzyme specificity was set to trypsin/P, allowing a maximum of two missed cleavages. Cysteine carbamidomethylation was set as fixed modification, while methionine oxidation and protein N-terminal acetylation were used as variable modifications. Global false discovery rate for both protein and peptides was set to 1%. The match-between-runs and re-quantify options were enabled. Intensity-based quantification options (iBAQ and LFQ) were calculated. Perseus software was used for data visualization (Tyanova et al, 2016); after canonical filtering (reverse, potential contaminants, and proteins only identified by site), only proteins with at least 1 unique peptide in all the replicates were considered as identified while only proteins with LFQ or SILAC ratio in all the replicates were defined as quantified. Pathway enrichment analysis was performed using the Metascape web software (92).

Survival probability plot was generated with UCSC Xena (http://xena.ucsc.edu), while log2 RNA seq data visualization was performed with firebrowse (http://firebrowse.org). Boxplot were generated with R studio, R (https://rstudio.com) or boxplotr web software (http://shiny.chemgrid.org/boxplotr/). STRING web software (https://string-db.org/) was adopted for functional protein association network visualization.

### Immunofluorescence in AID-DIvA cells

For immunofluorescence, around 1 × 10^4^ AID-DIvA cells (Aymard et al., 2014) were seeded in 24-well plates on glass coverslips in DMEM medium without antibiotics for 10h. Cells were then transfected with 10nM final concentration of control or target siRNAs using Lipofectamine™ RNAiMAX (Thermo Fisher Scientific), according to the manufacturer’s instructions, medium was then replaced with complete medium after 8h. 72h post-transfection, DSBs were induced with 300nM hydroxyl tamoxifen (4-OHT Sigma Aldrich T5648) for 4h. Repair was promoted *via* adding 500µg/mL Auxin (Indole-3-acetic acid sodium salt, Sigma Aldrich I5148) for 1h, triggering degradation of the AsiSI enzyme. Cells were fixed with 4% formaldehyde for 10min and permeabilized with 0.2% Triton X-100 for 10min while on shaking. For Extended Data Fig.5A, B, cells were subjected to click-chemistry reaction with 20mM of Cy5-or Biotin-azide (Jena Bioscience) for 30min. Biotin-azide samples were incubated afterwards with DNA hydrolysis buffer (1.5M HCl) for 30min, followed by three washes with PBS 1x. For figure 3D, U2OS cells were used in immunofluorescence and DSB were induced with 5Gy ionizing radiations with a Gammacell 40 Exactor (Best Theratronics) and cells were fixed and permeabilized 1h upon IR. All cells were incubated with blocking solution (1%BSA, 22.52mg/mL Glycine in PBS-Tween20 0.1%) for 1h while shaking. Immunostaining was performed overnight in a humidified chamber with antibody diluted in blocking solution. Antibodies against γH2AX (Millipore, 05-636), THRAP3 (Novus Biologicals, NB100-40848) were used at 1:1000 dilution, antibody anti-RAD51 (Millipore, PC-130) and anti-BRCA1 (Santa Cruz Biotechnology, sc6954) were incubated at 1:100 and 1:50, respectively. After washes, secondary antibodies were added at 1:500 dilution for 1h, followed by washes with PBS1x. In the second wash, Hoechst at 1:1000 was added. Upon coverslip mounting, images were acquired with a Zeiss Cell Observer inverted microscope (Zeiss) with an oil objective at 63x magnification. Images were analyzed with ImageJ software (imagej.nih.gov/ij), where Hoechst or DAPI was used to count the number of cells and define nuclei boundaries as ROIs. RAD51-, BRCA- and γH2AX-*foci* were counted within each nucleus with an in-house developed Java Macro after background subtraction with rolling ball radius of 50pixels. Minimum size restrictions were adopted and only *foci* with at least 0.2 and 0.15 micron^2 were counted for RAD51, BRCA and γH2AX, respectively. In all conditions, at least 50 cells were imaged and the number of *foci* was represented as boxplot in comparison with non-targeting silencing control. Significance was calculated with One-way ANOVA statistics and *, ** and *** correspond to p-value lower than 0.05, 0.001, and 0.0001, respectively. For co-localization between γH2AX and THRAP3 in figure 3D upper panel, “Colocalization threshold” ImageJ plug-in in was used adopting default settings.

### Proximity ligation assay

Proximity ligation assay was performed with Duolink® Proximity Ligation Assay (Sigma Aldrich-Merck, DUO92102) according to manufacturer’s instructions. In brief, 1 × 10^4^ U2OS cells where seeded on glass coverslip and after 12h were either left untreated, or DSBs were induced with 5Gy ionizing radiation with a Gammacell 40 Exactor (Best Theratronics). Cells were then fixed, permeabilized as in the immunofluorescence protocol above. Blocking and incubation with primary antibodies against γH2AX (Millipore, 05-636), THRAP3 (Novus Biologicals, NB100-40848) was performed overnight in a humidified chamber. PLA mouse and rabbit probes were added and ligated, before rolling circle amplification according to the manufacturer’s instructions. Slide preparation and imaging acquisition were performed as in the immufluorescence protocol above. DAPI staining was used to count nuclei and for defining nuclei boundaries as ROIs. PLA products per nucleus were counted with ImageJ using in-house developed Java Macro after background subtraction with rolling ball radius of 10pixels. Number of colocalization events are reported in boxplot for untreated and IR condition. Significance was calculated with One-way ANOVA statistics and *** correspond to p-value lower than 0.0001.

### Cell cycle analysis by flow cytometry

Flow cytometric analysis was performed as previously described (93). Briefly, cells were transfected with 10nM final concentration of control or target siRNAs using Lipofectamine™ RNAiMAX (Thermo Fisher Scientific), according to the manufacturer’s instructions. After 72h cells were fixed overnight with ice-cold 70% ethanol, and then permeabilized in phosphate buffer solution (PBS) containing 0.25% Triton X-100 (Sigma). DNA was stained with 50µg/ml propidium iodide (PI, Sigma-Aldrich) in PBS containing 0.1% Triton-X-100 and 200µg/ml DNase free RNase A (Sigma-Aldrich). Measurements were performed on a BD LSRFortessa flow cytometer (BD Biosciences) with FACSDiva software version 8.0.1 (BD Biosciences). 100.000 PI+ events were recorded for each condition from three independent experiments. Data analysis was performed using FlowJo X 10.0.7 software (FlowJo).

### Traffic light reporter (TLR) assay

TLR assay was performed as previously described (29, 79). In brief, U2OS-TLR cells were transfected with 10nM final concentration of control or target siRNAs using Lipofectamine™ RNAiMAX (Thermo Fisher Scientific), according to the manufacturer’s instructions. After 10h, cells were co-transfected with plasmids expressing I-SceI nuclease fused to infrared fluorescent protein (IFP) and donor plasmid expressing GFP donor sequence fused to blue fluorescent protein (BFP), using Polyjet™ in vitro transfection reagent (SignaGen Laboratories) according to the manufacturer’s instructions. Seventy-two hours after siRNA transfection, cells were harvested, and GFP and mCherry signals (reflecting HDR and NHEJ, respectively) were measured by four-color fluorescent flow-cytometry using a BD LSRFortessa flow cytometer (BD Biosciences). A minimum of 10.000 double-positive (IFP+/ BFP+) cells were recorded for each condition from three independent experiments. Data analysis was performed using FlowJo X 10.0.7 software (FlowJo). Results of siRNA-transfected cells were normalized to control siRNA-transfected cells. U2OS ID3-GFP (kindly provided by Dr. Ali Bakr, DKFZ) and U2OS m53BP1-mCherry cells (94) (kindly provided by Prof. Randi Syljuåsen, Oslo University Hospital) were used for compensation in flow cytometry.

### Colony formation assay, PARP inhibition and etoposide treatment

Colonigenic assays were performed as previously described (95) with some modifications. In brief, U2OS cells were plated overnight in complete medium without antibiotics at 20-25% density. Cells were then subjected to siRNA transfection with 10nM final concentration of control or target siRNAs using Lipofectamine™ RNAiMAX (Thermo Fisher Scientific), according with manufacturer’s instructions. After 24h post transfection, cells were collected, counted and seeded in a 6-well plate at a density of 1000 cells per well. 48h post transfection, cells were incubated for 2h with 1µM PARP inhibitor or solvent control; followed by DSB induction *via* 1µM etoposide treatment for 2h or solvent control. For figure 2E, cells were treated for 24h with etoposide at 0.1, 0.5, 1 or 10µM. For G9A colony formation assay, cells were exposed for 4h to 5µM G9Ai before incubation with PARPi. After 10-14 days, colonies were fixed for 10min in 70% ethanol, stained with crystal violet, destained in water and visualized. Colonies were counted with ImageJ software (imagej.nih.gov/ij) with an in-house developed Java Macro upon setting the image threshold and defining well boundary as ROI. Number of colonies or superficial area (normalized on reference) was normalized on non-targeting silencing control. Values from three biological replicates were averaged and displayed as mean ± standard deviation.

### Quantification and statistical analysis

SigmaPlot software was used to create graphs, perform statistical test and calculate p-values among at least three biological replicates. Unless stated otherwise, one-way ANOVA statistics was used for multiple comparison analysis. Each figure legend and respective method sections indicate both statistical significance and reference used for calculation of the p-value. For DDR-induced protein modulation, targets were classified as recruited or evicted if they fall in ±5%, respectively, of the 90 percentile log2 ratio distribution. Ratios of identified proteins with an intensity value (LFQ or iBAQ) only in either DDR-treated or untreated SILAC channel were replaced by a fixed value corresponding to ±6.67 in log2, respectively. Volcano plots were generated with Perseus software *via* a two-side t-test statistics, FDR> 0.05, S0 equal to 0.1. When reported, z-score normalization was performed per experiment (or column z-score normalization) with Perseus software. For the analysis of hPTM, the extracted ion chromatography (or XIC) was used as a measure of abundance of each modification. The relative abundance (or RA) is calculated as the XIC of a modified peak over the sum of all the XICs of all peaks for the same peptide multiplied by 100. In the heatmap RA over untreated time point are reported. The relative enrichment of a particular modification is calculated as the ratio of its RA in the ChIP over its RA in the chromatin input. Histone modifications with a log2 ratio higher or lower than 1 where considered as enriched or depleted, respectively.

## Notes

https://www.chromatin-proteomics.dkfz.de

### Competing Interest Statement

The authors have declared no competing interest.

### Summary of Updates

Further in-depth analyses as well as validation experiments have been performed. Moreover, to facilitate further exploration of chromatin dynamics during the DDR, we make our data available as a resource at www.chromatin-proteomics.dkfz.de, where identified proteins and histone modifications can be searched and displayed in the chromatin layers where they are quantified. Moreover, a dedicated tab shows dynamic trends of associated chromatin determinants (i.e. proteins and histone modifications), thus suggesting potential functional associations.

