## Extended data Figure for "Multi-layered chromatin proteomics identifies cell vulnerabilities in DNA repair"

Sigismondo et al. Extended Data Figure 1

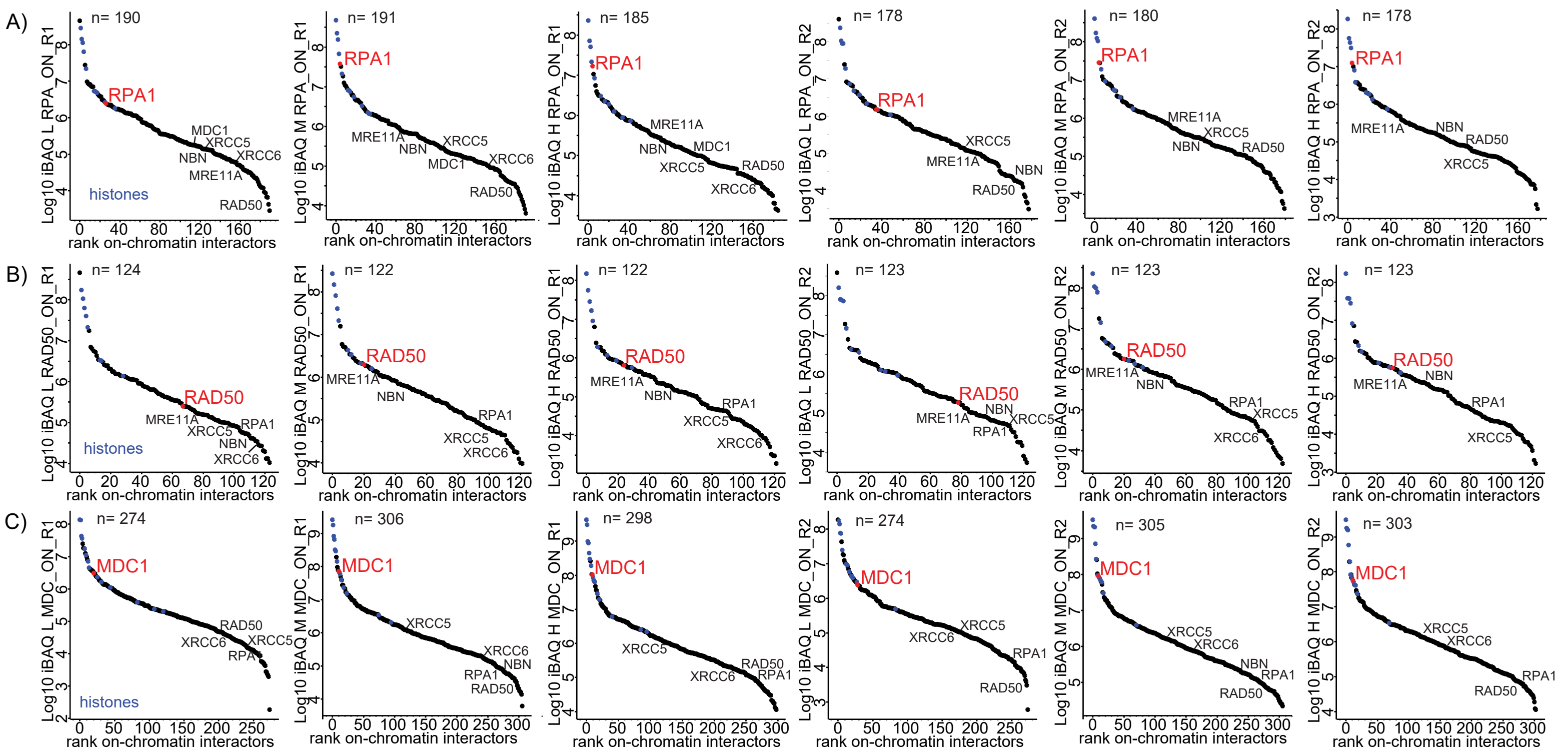


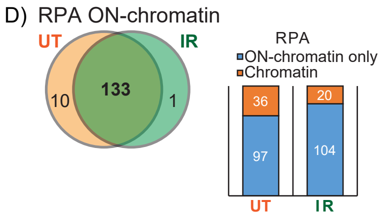


| RPA_ON-chromatin_Ionizing radiations-only | | |
|---|---|---|
| Protein IDs | Protein names | Gene names |
| V9HWF5 | Peptidyl-prolyl cis-trans isomerase A | PPIA |

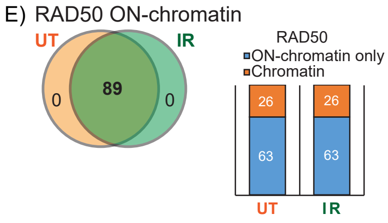


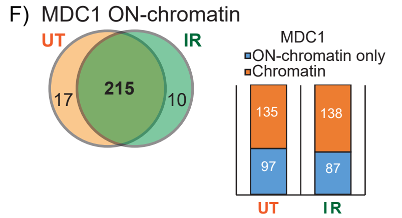


| MDC1_ON-chromatin_Ionizing radiation-only | | |
|---|---|---|
| Protein IDs | Protein names | Gene names |
| I6L9H2 | Cytosine-specific methyltransferase | DNMT1 |
| E5RFI6 | Histone deacetylase 2 | HDAC2 |
| F5H2U8 | High mobility group protein HMGI-C | HMGA2 |
| B3KSL5 | Putative RNA-binding protein Luc7-like 2 | LUC7L2 |
| P52701-2 | DNA mismatch repair protein Msh6 | MSH6 |
| O75475 | PC4 and SFRS1-interacting protein | PSIP1 |
| E7ESD9 | DNA repair protein RAD50 | RAD50 |
| A7E2E1 | Transcription activator BRG1 | SMARCA4 |
| Q86VX4 | Structural maintenance of chromosomes protein 3 | SMC3 |
| P49750-3 | YLP motif-containing protein 1 | YLPM1 |

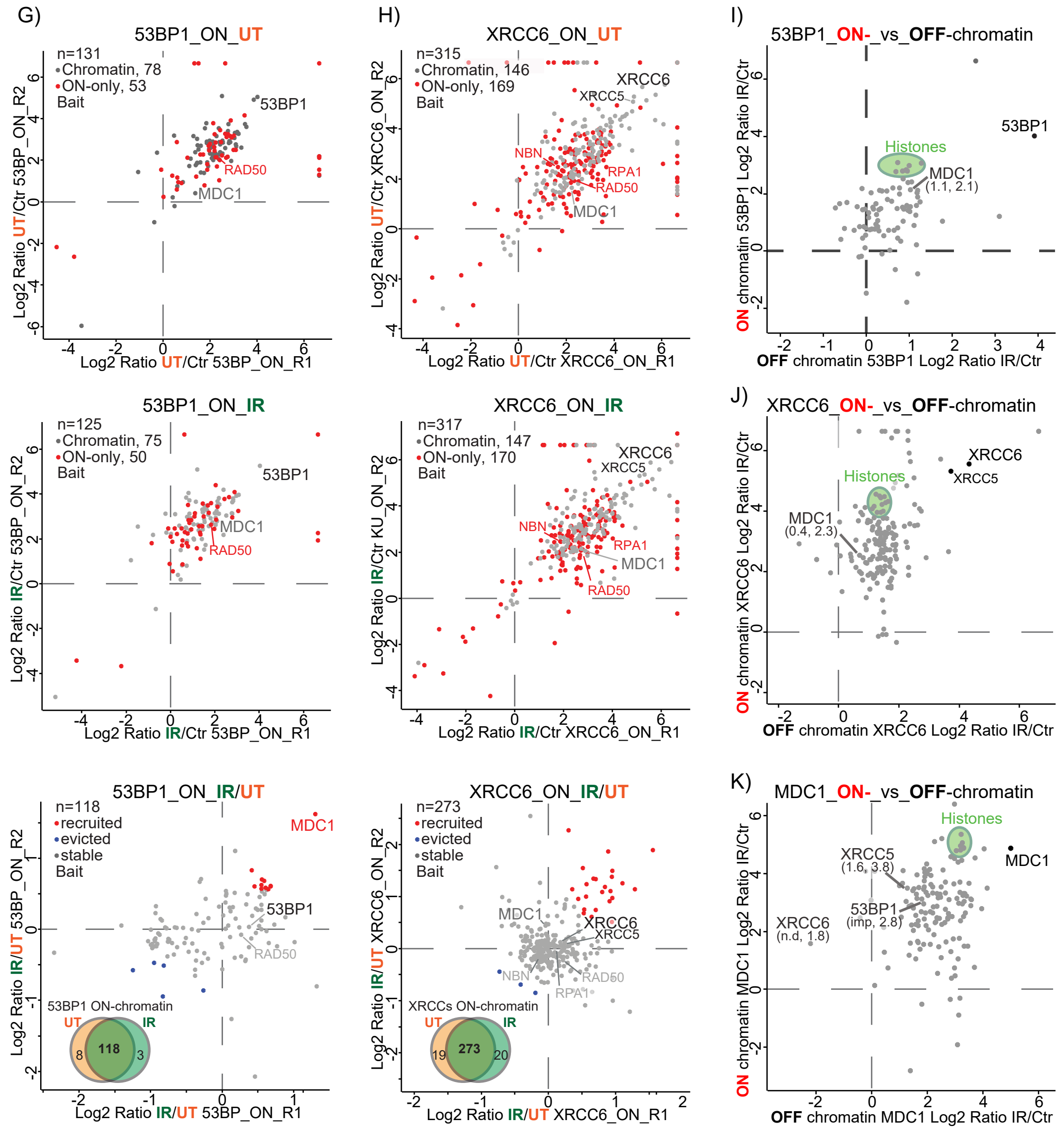

Sigismondo et al. Extended Data Figure 2

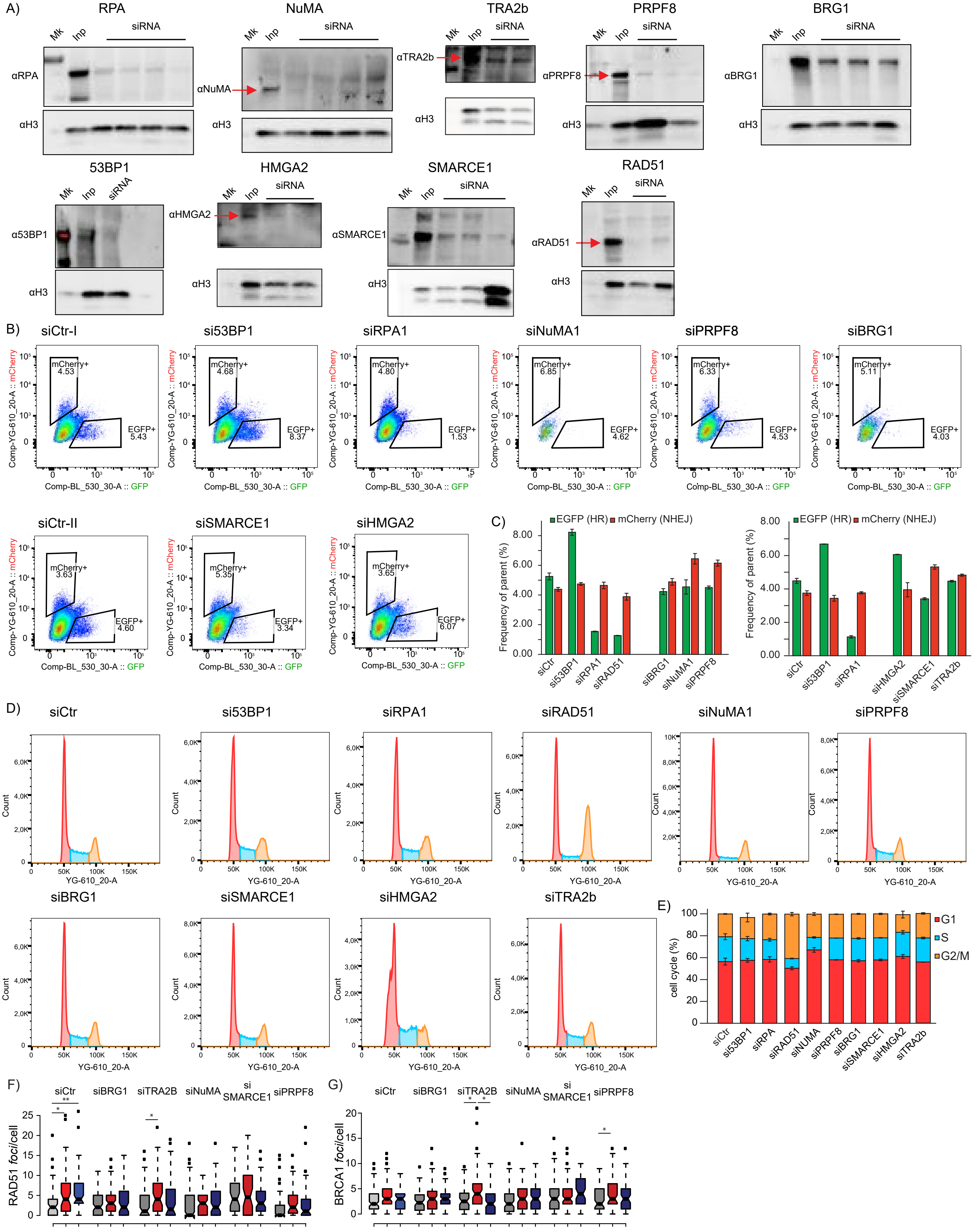

Sigismondo et al. Extended Data Figure 3

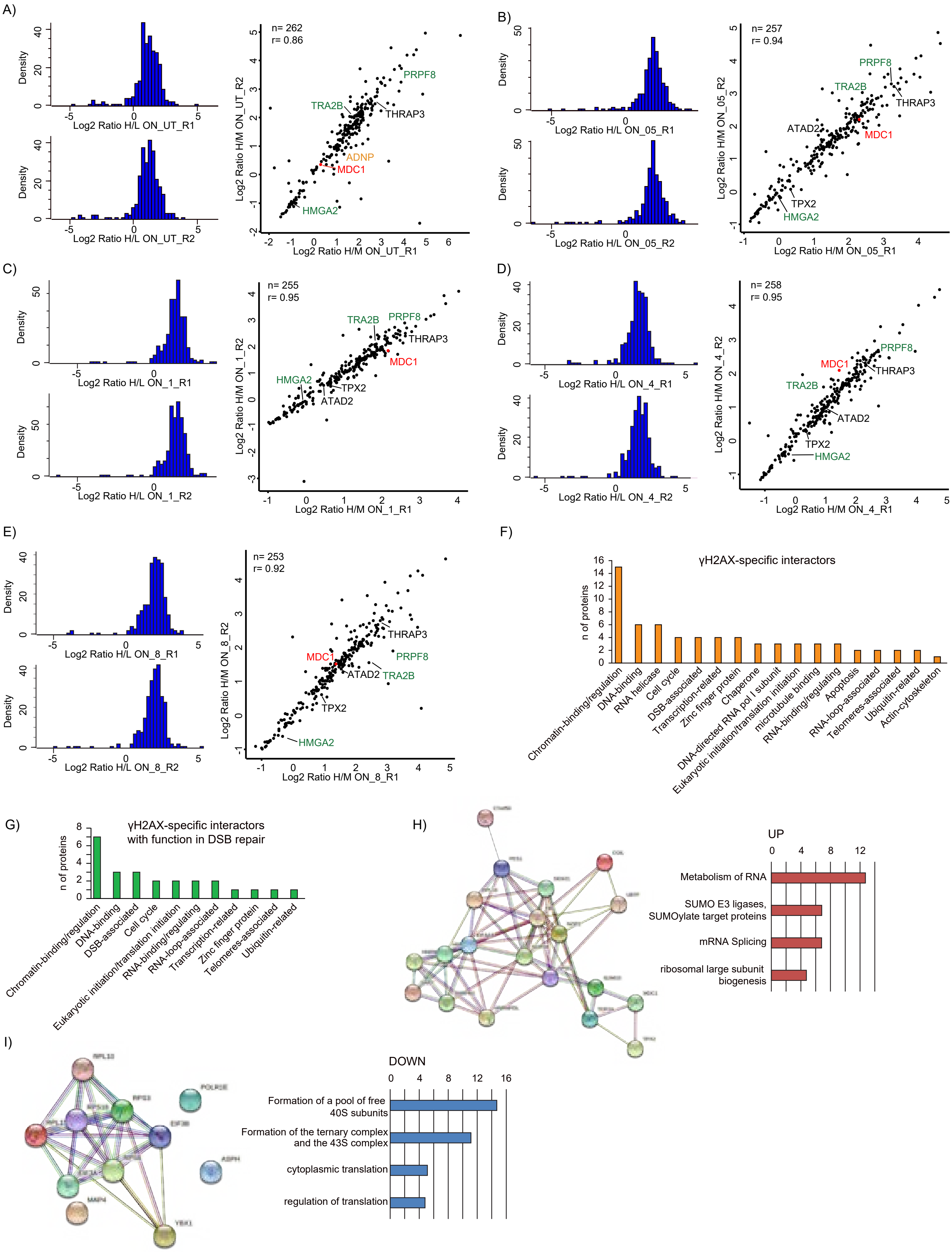

Sigismondo et al. Extended Data Figure 4

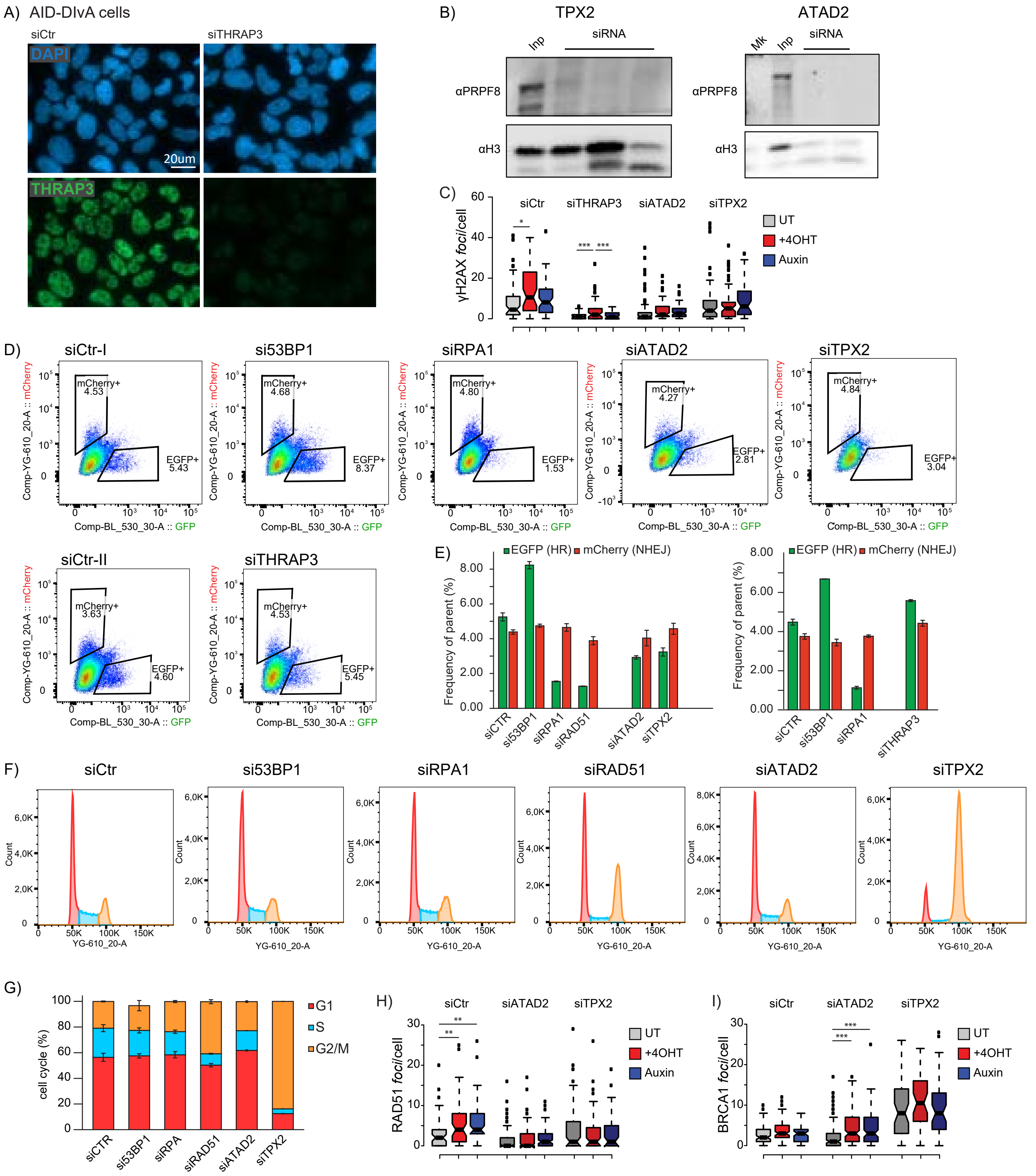

### Sigismondo et al. Extended Data Figure 5

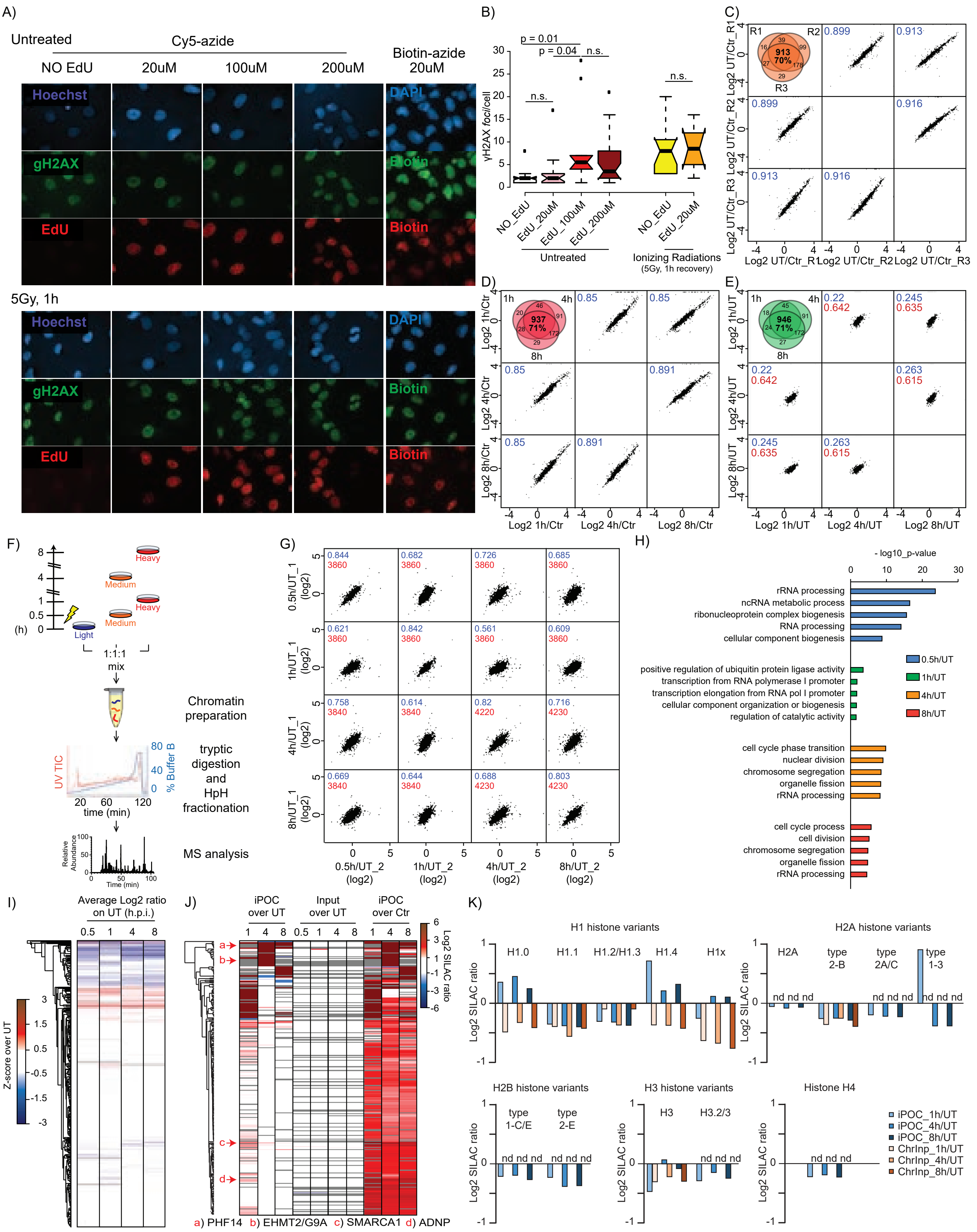

Sigismondo et al. Extended Data Figure 5 L-S

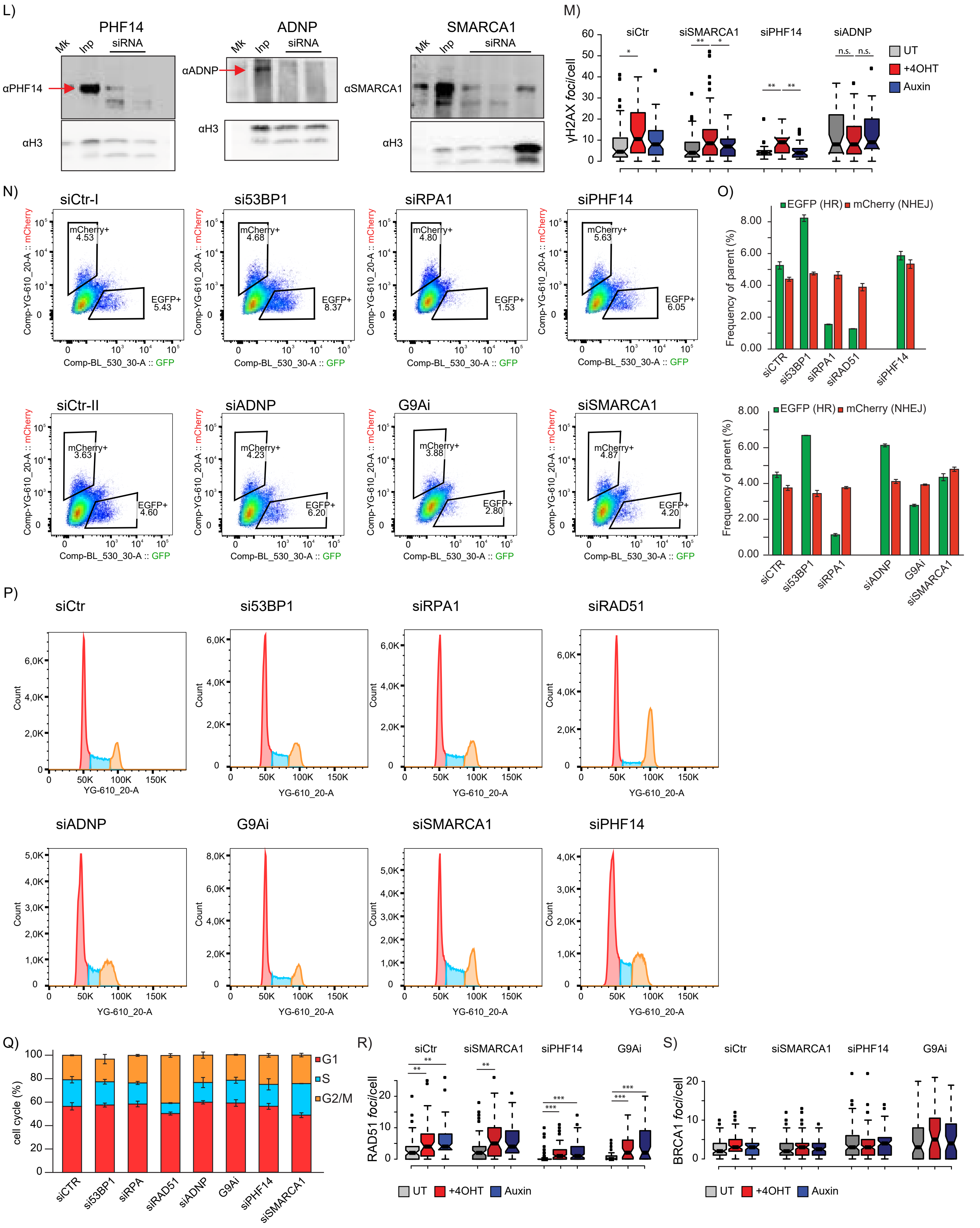

Sigismondo et al. Extended Data Figure 6

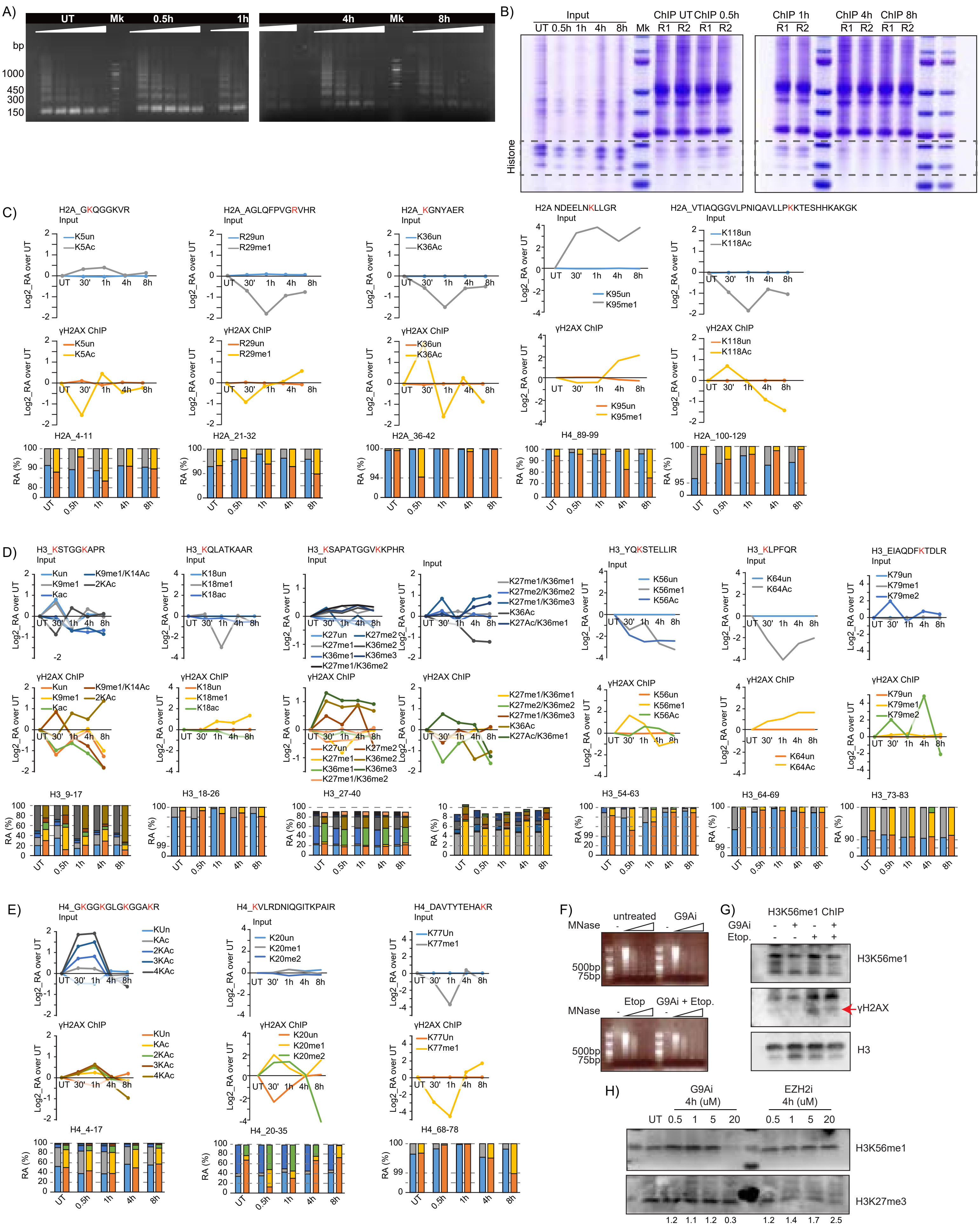
